# Benchmarking Oxford Nanopore Read Alignment-Based Structural Variant Detection Tools in Crop Plant Genomes

**DOI:** 10.1101/2022.09.23.508909

**Authors:** Gözde Yildiz, Silvia F. Zanini, Nazanin P Afsharyan, Christian Obermeier, Rod J Snowdon, Agnieszka A. Golicz

## Abstract

Structural variations (SVs) are larger polymorphisms (>50 bp in length), which consist of insertions, deletions, inversions, duplications, and translocations. They can have a strong impact on agronomical traits and play an important role in environmental adaptation. The development of long-read sequencing technologies, including Oxford Nanopore, allows for comprehensive SV discovery and characterization even in complex polyploid crop genomes. However, many of the SV discovery pipeline benchmarks do not include complex plant genome datasets. In this study, we benchmarked popular long-read alignment-based SV detection tools for crop plant genomes. We used real and simulated Oxford Nanopore reads for two crops, allotetraploid *Brassica napus* (oilseed rape) and diploid *Solanum lycopersicum* (tomato), and evaluated several read aligners and SV callers across 5×, 10×, and 20× coverages typically used in re-sequencing studies. Our benchmarks provide a useful guide for designing Oxford Nanopore re-sequencing projects and SV discovery pipelines for crop plants.

## 1. INTRODUCTION

Structural variations (SVs) are a major type of polymorphisms, which consist of insertions, deletions, inversions, duplications, and translocations. SVs are larger polymorphisms (>50 bp) compared with single nucleotide polymorphisms (SNPs) and small indels (insertions and deletions). Copy number variations (CNVs) and presence/absence variations (PAVs) occur due to these genomic polymorphisms (Alkan et al., 2011; Sedlazeck et al., 2018a). Insertions and deletions can have a strong effect on crop traits and have been shown to play a role in domestication and environmental adaptation (Gill et al., 2021; Tao et al., 2019; Yildiz et al., 2022; Zanini et al., 2022; Żmieńko et al., 2014). Until recently, the lack of high-quality reference assemblies and the complex nature of often large, polyploid genomes made comprehensive SVs exploration challenging in crop genomic research (Meyers and Levin, 2006; Yuan et al., 2021).

Development of long-read sequencing technologies such as Oxford Nanopore Technologies (ONT) (Jain et al., 2016) and Pacific Bioscience (PacBio) (Roberts et al., 2013) provided new opportunities for comprehensive SV discovery in crop plants. The sequencing accuracy of these technologies is continuously improving. Currently, PacBio HiFi consensus reads exceed 99% accuracy (Wenger et al., 2019) while ONT R10.3 raw reads accuracy exceeds 95% (Delahaye and Nicolas, 2021). The reduction in error rates facilitates downstream applications, including the production of high-quality genome assemblies, and SV detection. ONT sequencing in particular is being adopted in crop plant research for large scale re-sequencing projects of tens to hundreds of individuals (Alonge et al., 2020; Chawla et al., 2021; Lemay et al., 2022; Vollrath et al., 2021; Zhang et al., 2022). Despite the constant decrease in sequencing error rate, long-read technologies require specialized computational approaches to take advantage of them efficiently.

The two main approaches for SV discovery are *de novo* assembly-based and read alignment-based. *De novo* assembly-based approaches assemble reads into longer contigs and identify SVs by aligning assemblies (Wenger et al., 2019). Read alignment-based approaches directly align reads to reference genomes to discover SVs. *De novo* assembly-based methods perform better at finding larger variants (tens to hundreds of kbp long; exceeding the length of individual reads) but require sufficient amount of data to produce high-quality assemblies, which leads to substantial increase in cost of the experiments for larger crop genomes. However, read alignment-based approaches can perform well even at modest sequencing depths of 5× to 10× and use less computational resources, but the discovered SVs are limited to differences with the reference genome which makes this approach more suitable for larger re-sequencing projects (Coster et al., 2021). Several algorithms were developed for SV discovery from long-reads including Sniffles (Sedlazeck et al., 2018b), NanoVar (Tham et al., 2019), SVIM (Heller and Vingron, 2019), cuteSV (Jiang et al., 2020), and dysgu (Cleal and Baird, 2022), which have been comprehensively reviewed recently (Mahmoud et al., 2019; Yuan et al., 2021). Additionally, several long-read aligners are available such as minimap2 (Li, 2018), NGMLR (Sedlazeck et al., 2018a), Vulcan (Fu et al., 2021), and lra (Ren and Chaisson, 2021). Considering the continued development and improvement in read-alignment and SV detection algorithms and multitude of their possible combinations, their combined performances in SV detection demand realistic and up-to-date benchmarks to guide the selection of SV discovery tools.

In this study, we hypothesized that certain combination(s) of read aligners and SV discovery software will have superior performance in datasets representing complex crop genomes. We used real and simulated ONT reads for two crop plant genomes and evaluated several mappers and SV callers across coverages including 5×, 10×, and 20× typically utilized in re-sequencing studies. We chose to perform benchmarking on allotetraploid *Brassica napus* (oilseed rape) and diploid *Solanum lycopersicum* (tomato) as these two species represent different ploidy, have different SV profiles, and were already studied using Oxford Nanopore Technology. Our benchmarks provide a useful guide for researchers designing Oxford Nanopore re-sequencing projects and those designing SV discovery pipelines.

## 2. MATERIALS AND METHODS

### 2.1 Read Aligners, SV Callers, and Benchmarking Datasets

The SV callers included in the study were selected using several criteria: (1) citation count (used as a proxy for popularity in the research community), (2) publication date and maintenance status (excluding older tools that were no longer maintained), (3) ability to detect both insertion and deletion SVs from ONT data. The benchmarking approach involved four long-read aligners, including minimap2 (Li, 2018), NGMLR (Sedlazeck et al., 2018a), lra (Ren and Chaisson, 2021), and Vulcan (Fu et al., 2021) as well as five SV calling software namely Sniffles (v2) (Sedlazeck et al., 2018b), NanoVar (Tham et al., 2019), SVIM (Heller and Vingron, 2019), cuteSV (Jiang et al., 2020), and dysgu (Cleal and Baird, 2022). All aligners and SV caller versions are provided in detail in (**Table S1**). Three simulated datasets (Sim_ONT_Bn1, Sim_ONT_Bn2, and Sim_ONT_Sl) and publicly available data, for *B. napus* and *S. lycopersicum* genomes, were used. The real-world datasets for whole genome Nanopore sequencing of *B. napus* cv. King 10 (accession number: SRR15731030) (Vollrath et al., 2021) and *S. lycopersicum* cv. M82 (accession number: SRR16966224) (Alonge et al., 2021) were downloaded from NCBI Sequencing Read Archive. The ONT reads were randomly subsampled to 5×, 10×, and 20× coverages using Rasusa (Hall, 2022) to test the effect of sequencing depth on SV discovery.

### 2.2 Simulated Dataset Generation

For three simulated datasets (workflow for all simulations is presented in (**Figure S1**), new haplotypes including SVs were generated, and synthetic ONT reads were simulated using VISOR v1.1 (Bolognini et al., 2020). For simulation one (Sim_ONT_Bn1) 20,000 genomic intervals (mean: 750 bp, SD: 500 bp) were randomly drawn from the *B. napus* genome (Express 617 v1). A subset of 10,000 were denoted as deletions. For the remaining 10,000, denoted as insertions, the genomic start coordinate was retained, while the sequences corresponding to the genomic intervals were extracted, randomly re-assigned to the coordinates, and served as insertion sequences at those coordinates (**Figure S1**).

Simulations two and three, denoted Sim_ONT_Bn2 and Sim_ONT_Sl, were designed to reflect SVs found in real-world datasets. For Sim_ONT_Bn2 the assembled *B. napus* genomes Express 617 v1 (Lee et al., 2020) and Westar (Song et al., 2020) were aligned using minimap2 v2.24. SVs were detected using SVIM-asm v1.0.2 (Heller and Vingron, 2020). To reduce the effect of using minimap2 for benchmarking dataset generation, the SV locations were shifted by a randomly selected number in the (−5000, 5000) interval. This changed the exact SV site while maintaining the realistic distribution of SV sizes and locations along the genome. A random subset of 10,000 insertions and 10,000 deletions was drawn from all SVs to create the benchmarking dataset. SNPs discovered from short reads using bcftools v1.15.1 were also included. The SVs and SNPs were provided to VISOR to generate new haplotypes, which in turn were used for Oxford Nanopore read simulation. Sim_ONT_Sl was generated using the same strategy as for Sim_ONT_Bn2 but designed to reflect SVs of the *S. lycopersicum* genome. Heinz 1706 (Slycopersicum_691_SL4.0) and M82 (Alonge et al., 2021) assemblies were used for whole genome alignments. Due to smaller number of SVs, a random subset of 2,500 insertions and 2,500 deletions were drawn from all SVs.

To test the effect of sequencing depth on SV discovery, the datasets were simulated at 5×, 10×, and 20× coverage. The simulations provided the objective truth sets, which could be used to calculate SV precision, recall, and combined F1-scores. Precision describes the proportion of correct positive predictions among all positive predictions. It is calculated by dividing the true positives by overall positives. Recall describes the proportion of positive predictions made out of all positive elements in the dataset. It is calculated by dividing true positives by total number of relevant elements. F1-score combines precision and recall by taking their harmonic mean. Its value ranges from 0 to 1. F1-score close to 1 indicates high precision and recall. Using two different strategies for generating simulated datasets will make it possible to minimize analytical bias. If the same combination of tools performed best on all simulated datasets, this will likely reflect true superior performance.

### 2.3 Comparative Analyses

Express 617 v1 for the *B. napus* (Lee et al., 2020) and Slycopersicum_691_SL4.0 for the *S. lycopersicum* (Hosmani et al., 2019) were used as reference sequences. Simulated datasets and real subsampled reads at each coverage depth were aligned to respective reference genomes. The SV call sets were filtered using the following criteria: (1) number of minimum supporting reads: 5×: 3, 10×: 5, and 20×: 8, (2) SV type: INS or DEL (the most abundant SVs supported by all the benchmarked tools), (3) minimum SV length: 50 bp, (4) SV quality: SVs flagged as “PASS” (5) genotype: homozygous genotype for alternative allele (‘ 1/1’). For simulated data, precision, recall, and F1-scores of the SVs were computed for each combination of coverage depth, read aligner, and SV caller using Truvari v3.0.0 (English et al., 2022). Comparisons between results from the same tool combination across different coverages and different tool combinations across the same coverages were performed using surpyvor v0.8.1 (Jeffares et al., 2017). For real datasets, where no truth sets were available, we focused on within-dataset comparisons and how those compared to the results from simulated data. For each coverage, 20 different read aligner/SV caller combinations were used which led to a total of 60 different combinations for three coverages. All the relevant commands for simulated data generation and SV discovery are available in the **Supplementary Note.**

## 3. RESULTS

### 3.1 Selecting the Benchmarking Datasets

We chose to focus on two crop plant species *B. napus* (oilseed rape; genome size ~1.1 Gbp) and *S. lycopersicum* (tomato; genome size ~900 Mbp) because they are both important crops and their structural variation was previously studied using Oxford Nanopore Technologies (Alonge et al., 2020; Chawla et al., 2021). Whole Genome Alignment (WGA)-based SV discovery also suggested that they have quite different SV profiles with 38,666 SVs (Real_WGA_Bn, mean size: 2,068 bp, median size: 593 bp, 19,450 insertions and 19,216 deletions) discovered for *B. napus* and 7,108 SVs (Real_WGA_Sl, mean size: 3,029 bp, median size: 178 bp, 4,159 insertions and 2,949 deletions) discovered for *S. lycopersicum*.

Two simulated *B. napus* haplotypes (Sim_ONT_Bn1 and Sim_ONT_Bn2) and one simulated *S. lycopersicum* haplotype (Sim_ONT_Sl) were used to generate Oxford Nanopore reads at 5×, 10×, and 20× to test the effect of sequencing depth on SV discovery. The two publicly available real-world datasets, from *B. napus* (38×) and *S. lycopersicum* (68×), were subsampled with the same logic (Real_ONT_Bn, Real_ONT_Sl). The available graphical representation of a workflow for simulation and real data is shown in **Figure 1**.

**Figure 1:**
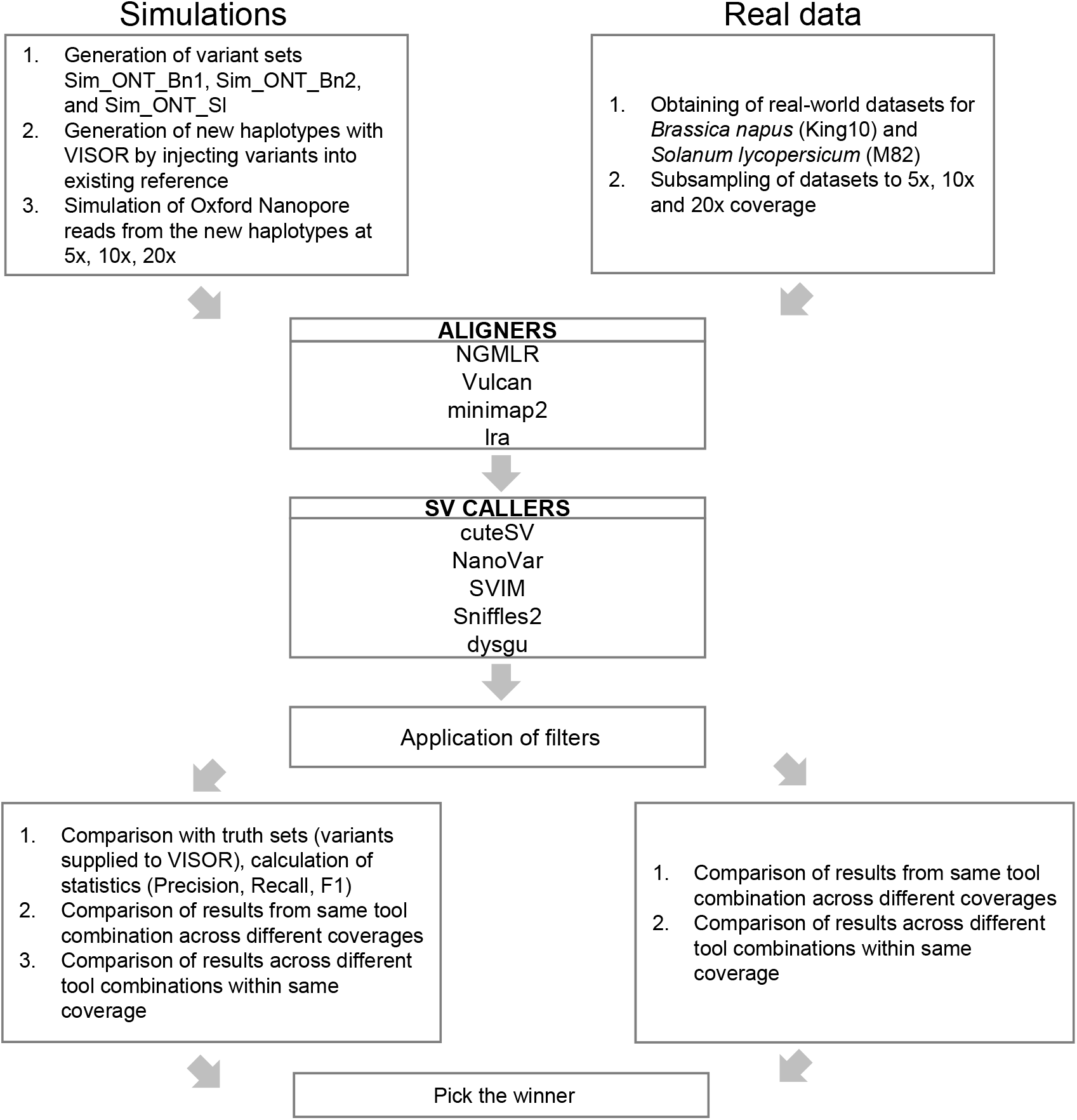
Graphical overview of the benchmarking workflow.

### 3.2 Characteristics of Structural Variant Truth Sets

The SVs supplied to VISOR to generate Sim_ONT_Bn1, Sim_ONT_Bn2, and Sim_ONT_Sl haplotypes served as three truth sets for our comparisons. The truth sets included deletions and insertions. The length distribution of truth set SVs is presented in **Figure 2**. Sim_ONT_Bn1 is unbiased in terms of the bioinformatics tools used, as the regions representing SVs were entirely randomly drawn from the *B. napus* genome. For any simulated dataset to reflect realistic SV distribution, SVs have to be discovered first and provided to the simulation software. Any relationship between tools used for SV identification for long-read dataset simulation and tools used for SV detection from these simulated reads (for example use of similar/same mapping algorithm) can result in inflated performance and biased results. However, Sim_ONT_Bn1 does not reflect realistic SV length and genomic distribution. To mitigate that Sim_ONT_Bn2 and Sim_ONT_Sl were created using SVs derived from real-world datasets. The two simulation strategies are complementary and should allow both unbiased and realistic assessment of SV calls. The median (mean) sizes (bp) for insertions and deletions were 800 (834) and 795 (825) for Sim_ONT_Bn1, 629 (1,959) and 594 (1,904) for Sim_ONT_Bn2 and 162 (3,178) and 165 (2,477) for Sim_ONT_Sl. Overall, the Sim_ONT_Bn2 and Sim_ONT_Sl truth sets had a wider range of insertion and deletion sizes. They were more reflective of true biological variation, making them more realistic than the Sim_ONT_Bn1 truth set.

**Figure 2:**
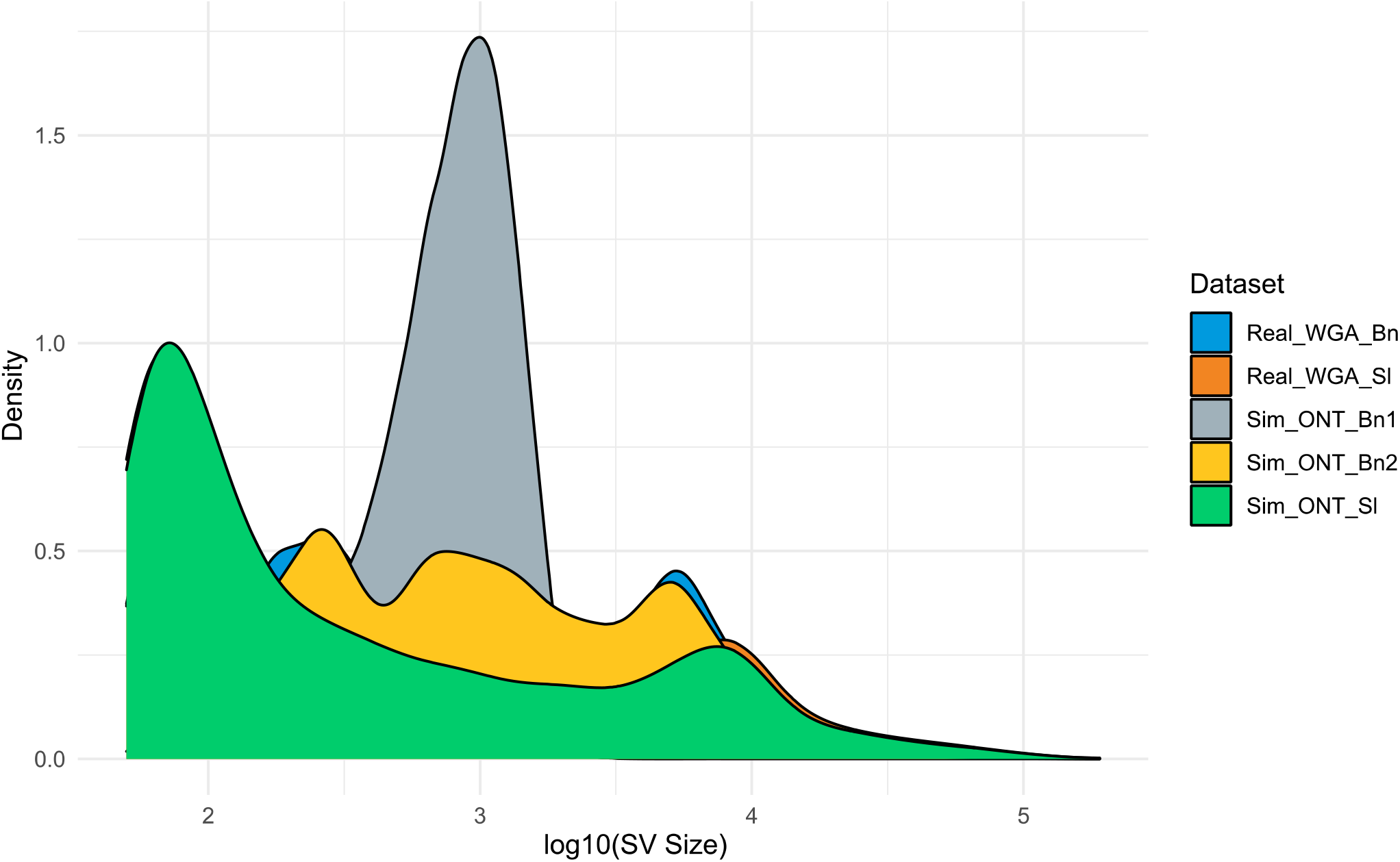
Size distribution of the real-world SV and SV from three benchmarking datasets.

### 3.3 Performance of Long Read Aligners

Subsampled *S. lycopersicum, B. napus*, and simulated reads were aligned using lra, minimap2, Vulcan, and NGMLR to the Slycopersicum_691_SL4.0, and Express 617 v1 reference genomes. Mapping statistics and run times of alignment against relevant reference genomes with different coverages of Sim_ONT_Bn1, Sim_ONT_Bn2, Sim_ONT_Sl, *B. napus* (Real_ONT_Bn), and *S. lycopersicum* (Real_ONT_Sl) real-world datasets are given in **Table S2**. Minimap2 had the shortest run time across all coverages. Conversely, NGMLR had the longest run time and also the lowest mapping rate. **Figure 3** shows mapping runtime (h:mm:ss or m:ss) for both simulation and real-world datasets with eight CPUs. Real_ONT_Bn dataset with 20× coverage was aligned ~220 hours by NGMLR and ~119 hours by Vulcan, compared to ~4 hours by minimap2 and ~5 hours by lra. Therefore, minimap2 and lra provided a greater speed advantage than NGMLR and Vulcan.

**Figure 3:**
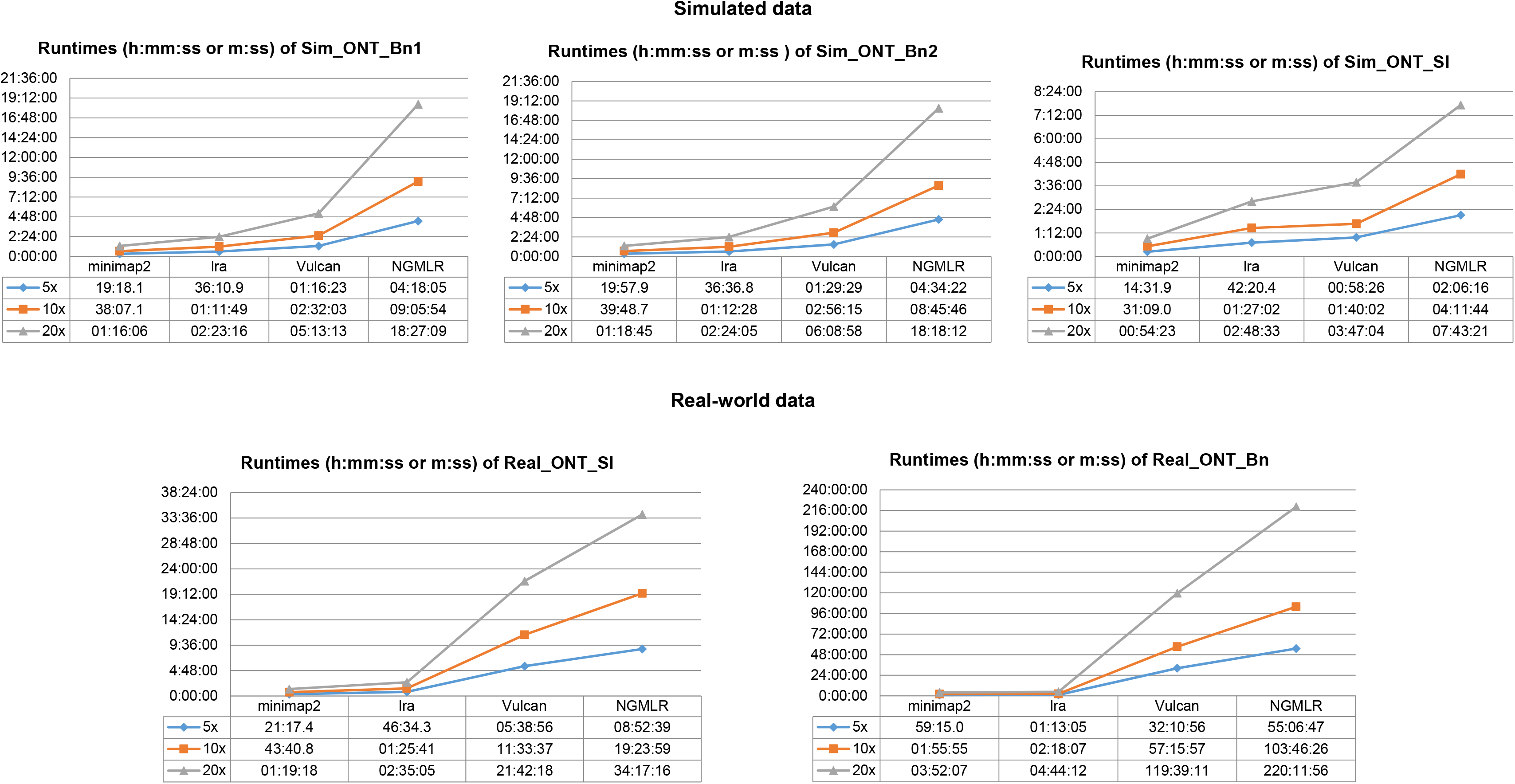
Read aligner run time (h:mm: ss or m: ss) for both simulation and real-world datasets with 5×, 10×, and 20× coverages (8 CPU). The reads were simulated with a mean length of 15,000 bp. The real-world datasets had a mean read length of 12,553 bp for *B. napus* and 22,339 bp for *S. lycopersicum*.

The run times increased with the higher coverages (**Figure 3**). Processing of real data took substantially longer than processing of simulated data. Moreover, Vulcan and minimap2 produced the highest proportion of mapped reads in Real_ONT_Bn (>96%), Real_ONT_Sl (96%-98%), and all simulated data (>98%) (**Table S2**). NGMLR reported the lowest proportion of mapped reads for Real_ONT_Bn (~81%) and Real_ONT_Sl (~76%), while lra and NGMLR resulted in similar statistics (96%-97%) for Sim_ONT_Bn1, Sim_ONT_Bn2, and Sim_ONT_Sl at each coverage. The combination of fast run time, good mapping rate, and the SV calling results presented below suggest that minimap2 is the top-performing aligner for simulated and real reads.

### 3.4 Performance of SV Callers on Simulated Data

#### 3.4.1 Performance using Sim_ONT_Bn1 as benchmark

We calculated the precision, recall, and F1-score of the SVs generated using different mapper and SV caller combinations using the Sim_ONT_Bn1 truth set. **Table S3** shows comparison of the precision, recall, and F1-scores for all mapper/SV caller combinations at the 5×, 10×, and 20× coverages. Each aligner/SV caller combination was evaluated with respect to total SVs, deletions, and insertions. **Figure 4** presents the corresponding F1-scores at 5× to 20× coverages. CuteSV after minimap2 alignment reached the highest F1-scores 5×:~0.90, 10×:~0.97, and 20×:~0.99 for total SVs, 5×:~0.91, 10×:~0.97, and 20×:~0.99 for deletions, and 5×:~0.89, 10×:~0.96, and 20×:~0.99 for insertions. At the lower end of coverage (5×), combination of minimap2/cuteSV provided a better advantage when compared to other mapper/SV caller combinations, especially in capturing insertions. Minimap2/Sniffles2 had second-best F1-scores (**Figure 4**). SVs detection by NanoVar was obtained directly from reads as NanoVar has its own internal mapping algorithm therefore the precision, recall, and F1-scores for different aligners are not included.

**Figure 4:**
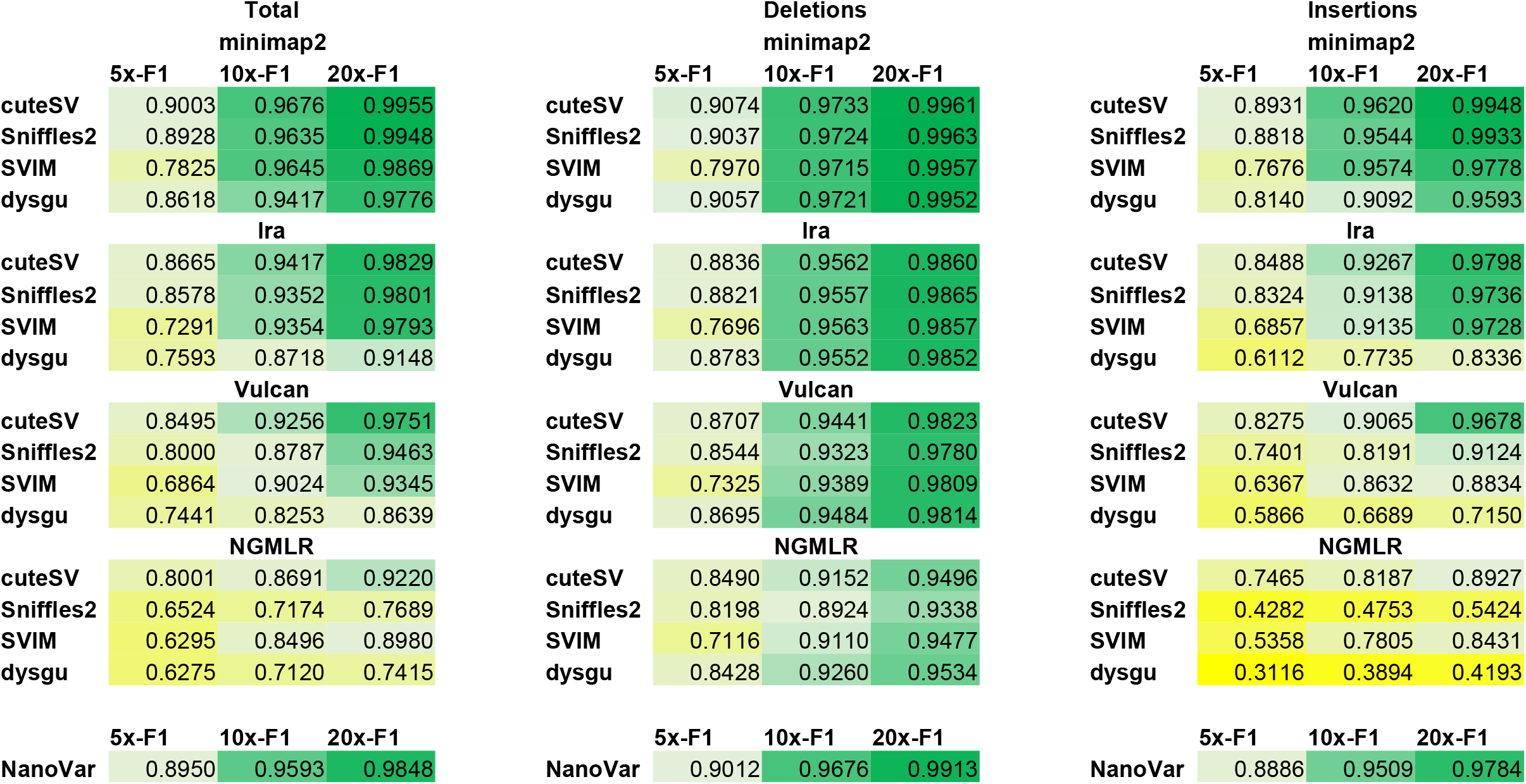
F1-scores of Sim_ONT_Bn1 including total SVs, deletions, and insertions at 5×, 10×, and 20× coverages for different combinations of read aligners and SV callers.

We also compared the total number of SVs, insertions, and deletions for all tested aligner/SV caller combinations. **Table S4** summarizes the number of SVs found at 5×, 10×, and 20× coverages. There were more discovered deletions than insertions regardless of coverage. The combinations of minimap2/cuteSV and minimap2/Sniffles2 detected the highest number of SVs at each coverage. We also analyzed how many of the SVs overlapped across different coverages while using the same tool combination and how many of the SVs overlapped across different tool combinations within the same coverage. **Data S1** shows the number of overlapping and unique SVs across coverages. Minimap2/cuteSV combination had the highest number of overlapping SVs. It also resulted in the highest proportion of overlapping SVs; 76.99% for all SVs, 79.19% for deletions, and 74.79% for insertions, while the minimap2/Sniffles2 combination (second best according to F1-scores) had the second highest percentage overlap; 75.35% for all SVs, 78.35% for deletions, and 72.33% for insertions **(Table S11** and **Figure 7)**. In addition, we performed comparisons across different tool combinations within the same coverage. **Data S2** displays the overlap, including the intersection sizes between SV calls and the Sim_ONT_Bn1 truth set. The highest number of overlapping SVs was found at 20x coverage, following minimap2 aligner. Our Sim_ONT_Bn1 results suggest that the combination of cuteSV and Sniffles2 with minimap2 alignment gave the best results achieving high F1-scores and capturing the highest number of overlapping SVs across coverages.

#### 3.4.2 Performance using Sim_ONT_Bn2 as benchmark

While Sim_ONT_Bn1 represents relatively short SVs randomly distributed along the genome, Sim_ONT_Bn2 reflects true biological variation in *B. napus*. **Table S5** presents comparison of the precision, recall, and F1-scores for all mapper/SV caller combinations at the 5×, 10×, and 20× coverages. **Figure 5** presents the F1-scores of SVs (total, insertions, and deletions) obtained using different combinations of aligners and variant callers across coverages. CuteSV following minimap2 alignment again was the top performing combination with the highest overall F1-score values 5×:~0.87, 10×:~0.93, and 20×:~0.96 for total SVs, 5×:~0.90, 10×:~0.96, and 20×:~0.98 for deletions, and 5×:~0.83, 10×:~0.89, and 20×:~0.94 for insertions. Especially, at low 5× coverage, this combination performed better than others. Minimap2/Sniffles2 had the second highest F1-scores at 20× coverage as in Sim_ONT_Bn1. However, minimap2/dysgu F1-score for insertions at 5× and 10× was higher than Sniffles2 after the minimap2 alignment.

**Figure 5:**
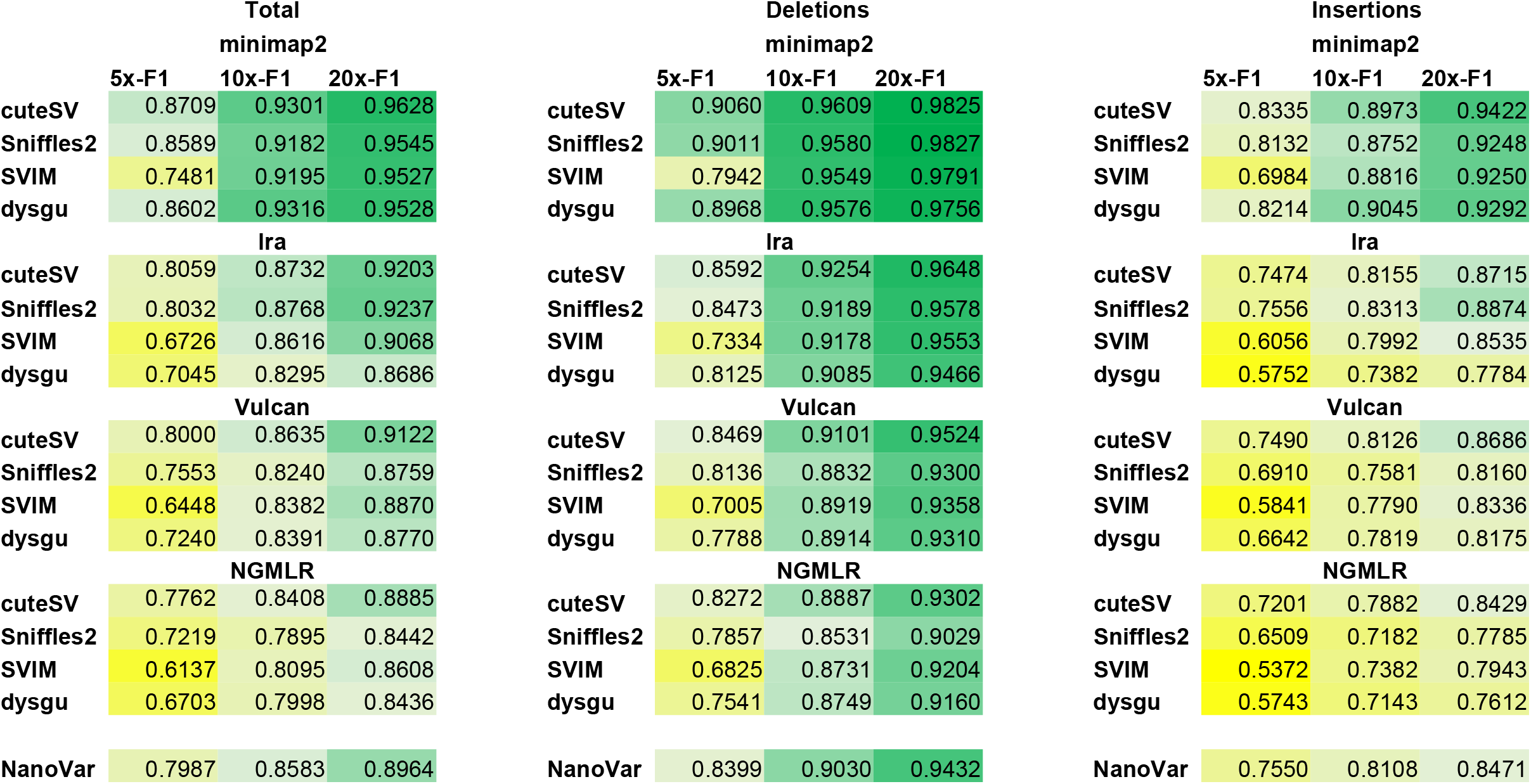
F1-scores of Sim_ONT_Bn2 including total SVs, deletions, and insertions at 5×, 10×, and 20× coverages for different combinations of read aligners and SV callers.

In addition, the total number of SVs, the total number of insertions, and deletions for all combinations of tested aligners and SV callers were compared. **Table S6** summarizes the total number of SVs detected at 5×, 10×, and 20× coverages. Minimap2/cuteSV found the highest number of SVs at each coverage like in Sim_ONT_Bn1. Again, more deletions than insertions were found for all aligner and SV caller combinations across different coverages. We also analyzed how many of the SVs overlapped across different coverages while using the same tool combination and how many of the SVs overlapped across different tool combinations within the same coverage. **Data S3** lists the number of overlapping SVs across different coverages using the same tool combination. Minimap2/cuteSV combination had the highest number of overlapping SVs. It also had the highest proportion of overlapping SVs; 73.95% for all SVs, 80.05% for deletions, and 67.44% for insertions. The minimap2/dysgu combination was second best detecting 73.23% for all SVs, and 67.28% for insertions. Minimap2/Sniffles2 combination was the second best for deletions with 79.14% overlap **(Table S11** and **Figure 7)**. **Data S4** displays overlap between results from different SV callers within the same coverage after each aligner, including the intersection with the Sim_ONT_Bn2 truth set. The highest number of overlapping SVs was found at 20x coverage, following minimap2 aligner. Overall, in Sim_ONT_Bn2, the combination of cuteSV after minimap2 alignment gave the best results both in terms of F1-Scores and concordance across coverages.

#### 3.4.3 Performance using Sim_ONT_Sl as benchmark

Sim_ONT_Sl represents the true biological variation of *S. lycopersicum*. **Table S7** presents comparison of the precision, recall, and F1-scores for all mapper/SV caller combinations at the 5×, 10×, and 20× coverages. **Figure 6** shows the F1-score of SVs (total, insertions, and deletions) identified using combinations of the different aligners and variant callers. CuteSV and Sniffles2 with minimap2 alignment were top performers with the highest F1-score values (5×:~0.85, 10×:~0.92, and 20×:~0.94) for total SVs, (5×:~0.88, 10×:~0.95, and 20×:~0.97) for deletions, and (5×:~0.81, 10×:~0.88, and 20×:~0.91) for insertions. Lra/Sniffles2 combination had the best F1-score for insertions for each coverage.

**Figure 6:**
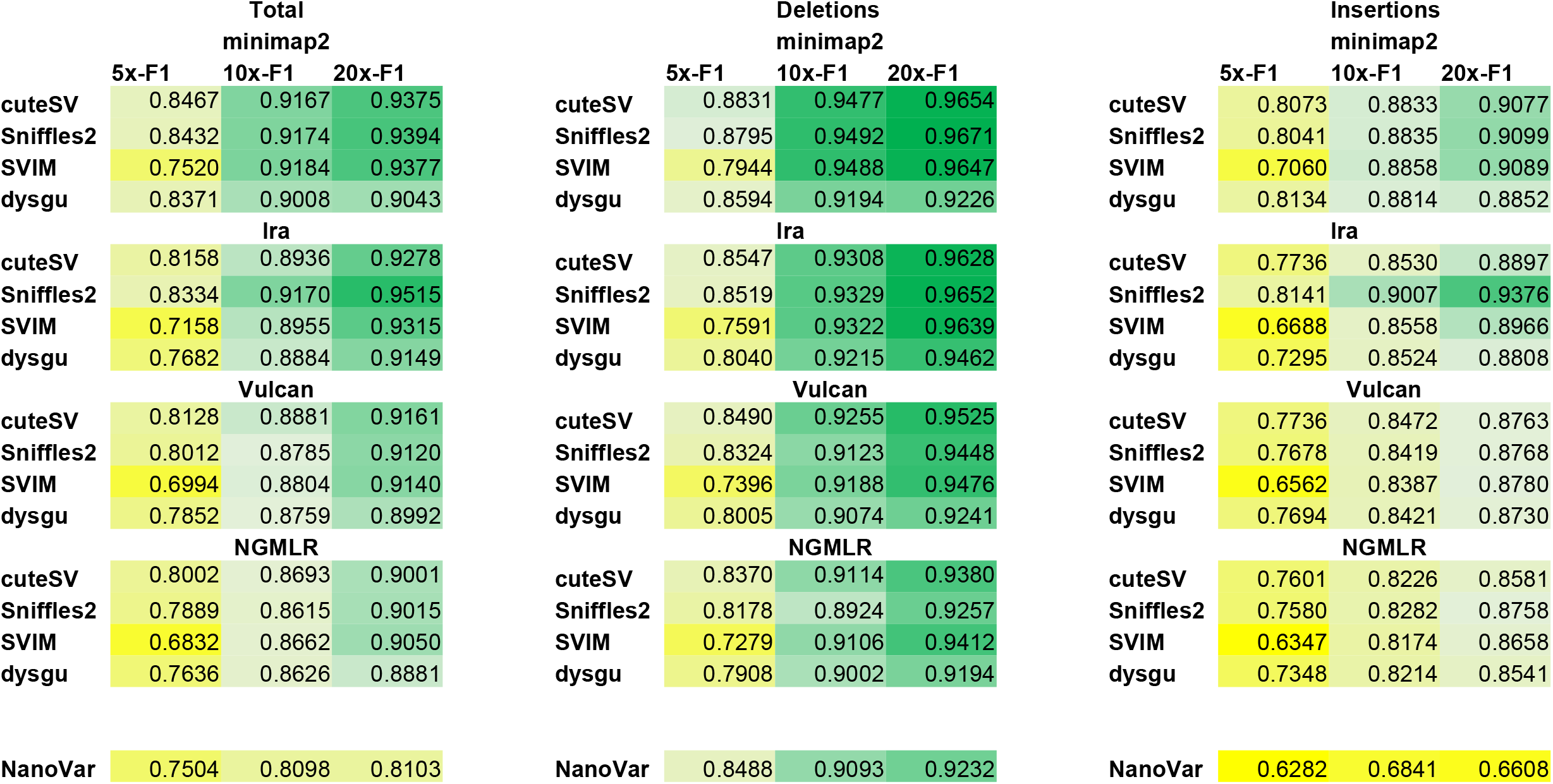
F1-scores of Sim_ONT_Sl including total SVs, deletions, and insertions at 5×, 10×, and 20× coverages for different combinations of read aligners and SV callers.

In addition, the total number of SVs, the total number of insertions, and deletions for all tested aligner/SV caller combinations were compared. **Table S8** summarizes the total number of SVs at 5×, 10×, and 20× coverages. Again, more deletions than insertions were found for all aligner and SV caller combinations across coverages like in the previous simulated datasets. The number of SVs overlapping across coverages while using the same tool combination and the number of SVs overlapping across different tool combinations but within the same coverage were also calculated. **Data S5** shows the number of overlapping SVs across different coverages using the same tool combination. Minimap2/dysgu combination had the highest number of overlapping SVs. However, minimap2/cuteSV combination found the highest proportion of overlap; 73.49% for all SVs, 77.52% for deletions, and 68.98% for insertions, while the minimap2/Sniffles2 combination was second best detecting 72.73% for all SVs, 76.32% for deletions, and 68.72% for insertions **(Table S11 and Figure 7)**. Although minimap2/dysgu found the highest number of SVs at each coverage in Sim_ONT_Sl, the proportion of overlapped SVs was reported as 68.82%. **Data S6** displays overlap between results from different SV callers within the same coverage after each aligner, including the intersection with Sim_ONT_Sl truth set. The highest number of overlapping SVs was found at 20x coverage, following minimap2 aligner. Overall, in Sim_ONT_Sl, the combination of cuteSV and Sniffles2 after minimap2 alignment gave the best results both in terms of F1-Scores and concordance across coverages.

**Figure 7:**
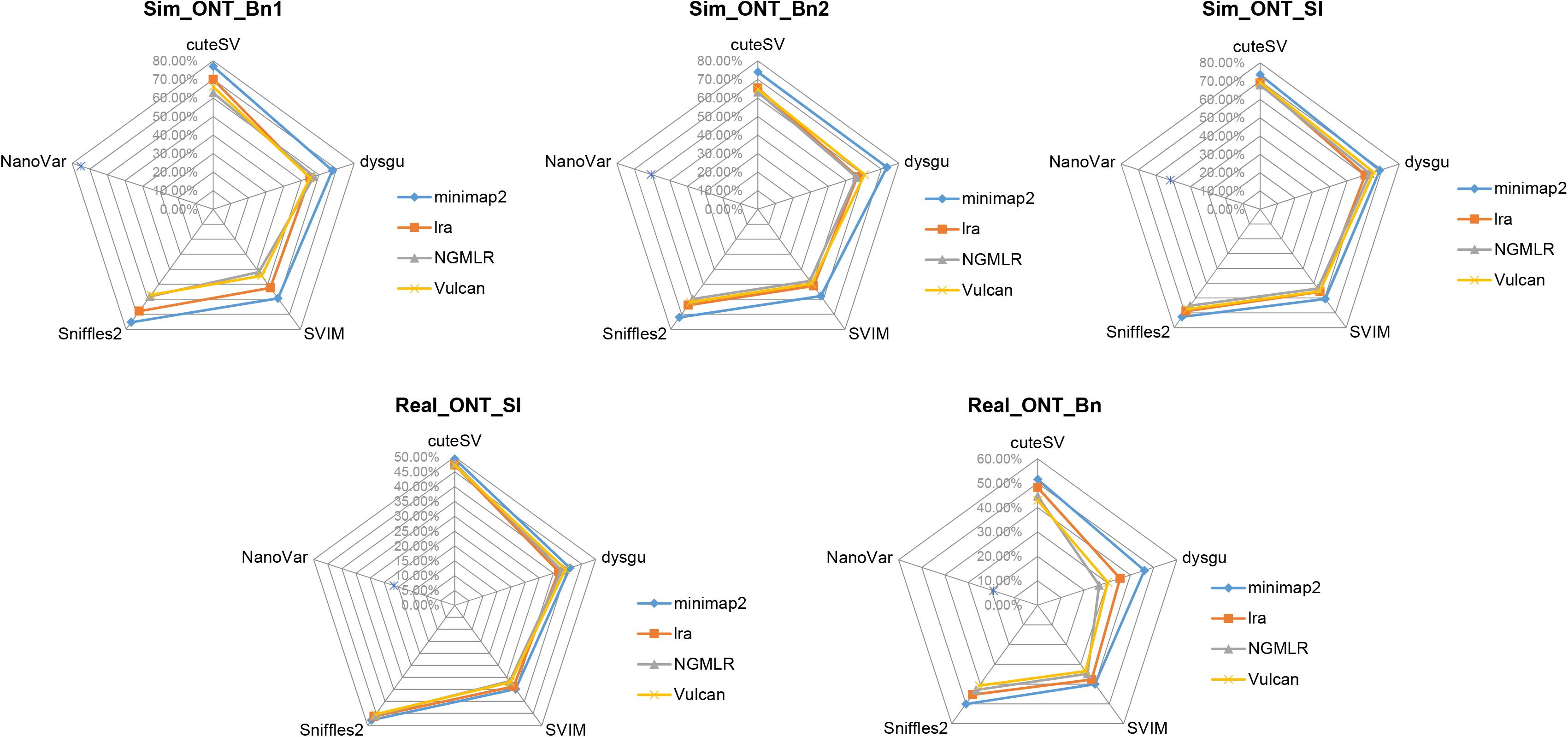
Proportion of overlapped SVs (%), across 5×, 10×, and 20× coverages for simulated and real-world datasets.

### 3.5 Performance of SV Callers on Real-World Data

While tool performance on simulated data provides a useful guide, real-world datasets usually provide additional unaccounted-for complexity and challenges. After finding the best combinations in simulated data, we investigated whether the pattern would be similar in real-world datasets. Since for the real-world data we do not have an objective truth set, they were only evaluated from two perspectives which are the congruence of results when using the same tool combination across different coverages and when using different tool combinations within the same coverage.

#### 3.5.1 Performance on *B. napus* Real-World ONT Data

*B. napus* ONT real dataset (Real_ONT_Bn) was evaluated using the above-described strategy. **Table S9** shows the number of SVs from all tested combinations at different coverages in *B. napus*. The minimap2/cuteSV and minimap2/dysgu combinations within all coverages captured the highest number of total SVs, deletions, and insertions. Overall, a higher number of deletions than insertions was detected for all aligner and SV caller combinations at different coverages. The number of overlapped SVs across coverages for the same SVs caller/aligner combinations was calculated (**Data S7**). Minimap2/cuteSV combination found the highest proportion of overlapping SVs discovered at different coverages using the same combination of tools (51.53% of total SVs, 54.52% of deletions, and 47.91% of insertions), while the minimap2/sniffles2 combination was second best, detecting overlap of 50.1% for all SVs, 54.56% for deletions, and 44.92% for insertions across coverages **(Table S12 and Figure 7)**. Although the minimap2/dysgu combination found more SVs, the percentage of intersecting SV was low. NanoVar detected the lowest proportion of overlapping SVs across coverages (19.04% of total SVs, 25.07% of deletions, and 10.21% of insertions) and discovered more unique SVs. Surprisingly we noticed a high proportion of heterozygous genotypes (0/1) in SV calling results for Real_ONT_Bn, considering that the data represented a highly inbred elite line (Vollrath et al., 2021). **Table S14** shows the number of SVs genotyped as homozygous and heterozygous in real-world data. As our SV filtering required the genotypes to be homozygous for the alternative allele (1/1) these heterozygous calls were removed prior to analysis. We also investigated the overlap in SV calls across different tool combinations within the same coverage (**Data S8**). We observed that a substantial proportion of deletions and insertions were shared by most SV callers, with the largest number of overlapping SVs at 20×, following minimap2 alignment.

#### 3.5.2 Performance on *S. lycopersicum* Real-World ONT Data

We performed a similar evaluation for the real-world dataset of *Solanum lycopersicum* (Real_ONT_Sl). **Table S10** shows the number of SVs found from all tested combinations at different coverages. The minimap2/dysgu combinations at 5×, 10×, and 20× captured the most SVs. Additionally, for *S. lycopersicum* all tool combinations with the exception of NanoVar found more insertions than deletions at each coverage. We also calculated the number of overlapping SVs while using the same tool combination across different coverages (**Data S9**). Minimap2/cuteSV combination found the highest proportion of overlapping SVs; 49.34% for all SVs, 49.63% for deletions, and 49.16% for insertions, while the minimap2/sniffles2 combination detected 47.80% for all SVs, 49.41% for deletions, and 46.61% for insertions. Even though the minimap2/dysgu combination found more SVs, the percentage of common SVs (40.82%) was low like Real_ONT_Bn data. NanoVar again detected the lowest proportion of overlapping SVs (21.57% for all SVs, 31.20% for deletions, and 12.16% for insertions), and it discovered more unique SVs like for the Real_ONT_Bn dataset **(Table S12 and Figure 7)**. Again, we also tested overlaps between SV calls within the same coverage, but across different tool combinations (**Data S10**). The largest number of overlapping SVs was found at 20×, following minimap2 alignment.

#### 3.5.3 The Unique Features of Real-World Datasets

We found a surprisingly high proportion of heterozygous calls in the real-world datasets given the highly inbred nature of the material used for sequencing. A high proportion of those is therefore likely SV discovery/genotyping errors. More heterozygous calls were found in the *B. napus* than the *S. lycopersicum* dataset. *B. napus* is an allotetraploid species, which undergoes reciprocal and non-reciprocal homeologous exchanges (HEs; exchanges of large corresponding chromosome segments between subgenomes). Non-reciprocal HEs could potentially cause erroneous SV calls if there are HE present in the reference, but absent in the sample. As a result, reads will have no corresponding mapping location and may be mis-mapped. To test such a scenario, we used the Sim_ONT_Bn2 dataset (20×, minimap2 for mapping, and cuteSV for SV detection) and two versions of the modified Express 617 reference. In the first version, we replaced chromosome A01 by C01 (two C01 chromosomes and no A01). In the second version, we replaced chromosome C01 by A01 (two A01 chromosomes and no C01). In both cases, the use of the modified reference resulted in an increased number of heterozygous (162.3% for reference with A01 missing, and 237.1% for reference with C01 missing), but not homozygous calls across all chromosomes (**Figure 8**), suggesting the non-reciprocal HEs can contribute to produce erroneous heterozygous calls.

**Figure 8:**
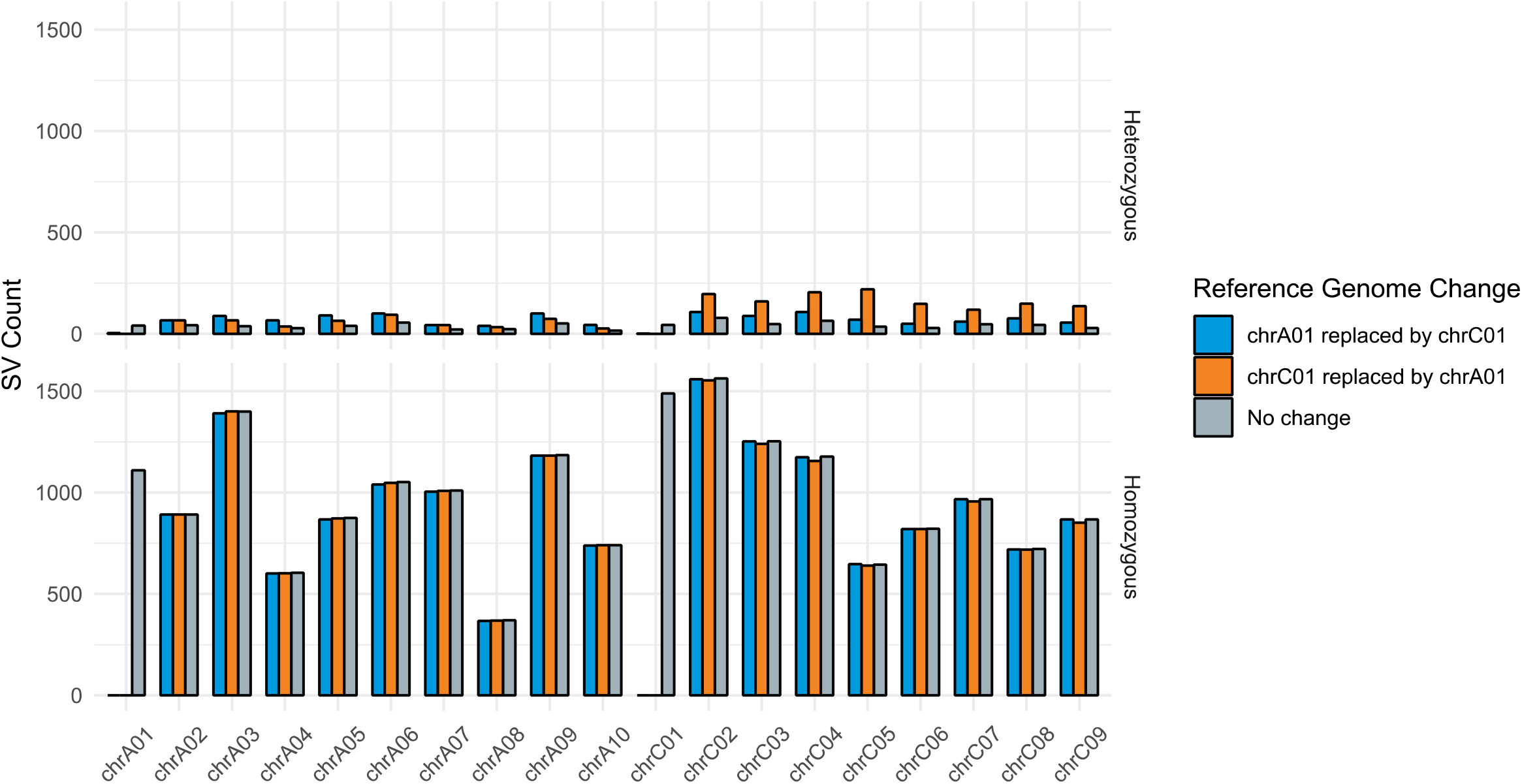
The effect of non-reciprocal homeologous exchanges on SV discovery. Non-reciprocal homeologous exchanges were simulated by replacing chromosome A01 by C01 and C01 by A01.

### 4. DISCUSSION

Many of the SV detection tools are benchmarked primarily on human/animal datasets, (Bolognini and Magi, 2021; Coster et al., 2019; Dierckxsens et al., 2021; Jiang et al., 2020; Jiang et al., 2021; Zhou et al., 2019), however the complexity and different SV profiles of crop plant genomes might bring unique challenges. Therefore, to guide the design of large-scale long-read re-sequencing studies, this study performed comprehensive benchmarking of popular SV calling tools with a focus on tool performance at lower sequencing coverage. For this purpose, we designed two data simulation strategies representing both unbiased and realistic benchmarking datasets reflecting structural variation for two major crops oilseed rape (*B. napus*) and tomato (*S. lycopersicum*).

Four long-read aligners (minimap2, NGMLR, lra, and Vulcan) and five SV callers (Sniffles2, SVIM, cuteSV, dysgu, and NanoVar) were tested to detect SVs, particularly deletions and insertions. Alignment time varied widely between the four aligners, while differences in the proportion of mapped reads were moderate. As expected, higher sequencing coverage and reference genome size length increased the run time of the mapping algorithms. The real-world datasets required more time at the same coverage and reference genome size, which most likely reflected additional complexity not captured in simulations. Overall, the results found minimap2 to be the best performing aligner for SV calling applications, which also had the fastest run time and the most mapped bases. Recent benchmarking studies on human data also recommended minimap2 among tested aligners such as GraphMap, LAST, and NGMLR (Bolognini and Magi, 2021; Coster et al., 2019; Zhou et al., 2019).

We found that similar tool combinations (especially cuteSV, followed closely by Sniffles2 and dysgu after minimap2 alignment) had superior performance across all the simulated datasets. The findings are in line with a recent study reporting that cuteSV performed better than other tested SV tools such as Sniffles1, SVIM, and pbsv for precision and recall at both SV calling and genotyping in human datasets (Bolognini and Magi, 2021). Increasing coverage improved recall and F1-scores for all tested SVs calling combinations, confirming that the probability of detecting quality SVs increases with more sequencing coverage (Jiang et al., 2021). However, even at low coverages (5×) using cuteSV, Sniffles2, and dysgu for SV detection from reads aligned by minimap2 achieved >0.8 F1-scores on simulated datasets, suggesting that Oxford Nanopore technology might be suitable for large-scale low coverage re-sequencing projects. While the lack of objective truth sets for real-world datasets precludes similar comparisons, the results revealed that tool combinations with best performance for simulated datasets also had the most consistent outcome across the range of coverages.

The criteria for filtering SV in this study were quite stringent, including retaining only SV genotyped as homozygous for alternative allele (1/1). While in simulated datasets the number of SV genotyped as heterozygous was relatively low, the proportion was much higher for real-world datasets, especially in *B. napus*. We found that in *B. napus*, the presence of homeologous exchanges will likely contribute to the erroneous discovery of heterozygous SV. *B. napus* is well known to harbour wide-spread non-reciprocal homeologous chromosomal exchanges even extending to whole chromosomes, e.g. for chromosomes A01 and C01 as simulated here (Udall et al., 2005). The finding underlies the importance of species-specific consideration when interpreting SV discovery results. The presence of HEs likely explains only a proportion of the observed heterozygous calls and other factors need to be considered as well, including other sources of mis-mappings, genotyping errors, and residual heterozygosity in samples.

In conclusion, we found that for homozygous/inbred genotypes often used in crop studies a substantial proportion of SVs can be discovered/genotyped at coverages as low as 5×, making Oxford Nanopore technology a suitable option for larger-scale re-sequencing studies. At this time, following our benchmarks we recommend using the minimap2/cuteSV combination as it achieves good precision and recall at SV calling and found the highest overlap between SVs across coverages. The performance of minimap2/cuteSV was followed closely by minimap2/Sniffles2 for both simulated and real datasets.

## DATA ACCESS

Supplementary Data and Files can be accessed under the following link: https://osf.io/9c5hz/

## ACKNOWLEDGMENTS

This work was supported by the Alexander von Humboldt Foundation in the framework of Sofja Kovalevskaja Award to AAG and the German Research Foundation (DFG) project number 458716530 to RJS.

## AUTHOR CONTRIBUTIONS

GY performed research, wrote the manuscript

SFZ assisted in the analysis, wrote the manuscript

NPA provided critical comments, wrote the manuscript

CO provided critical comments, wrote the manuscript

RJS edited the manuscript

AAG conceived research, supervised research, wrote the manuscript, acquired funding

## Notes

### Competing Interest Statement

The authors have declared no competing interest.

