## Supplementary figures and images for "Benchmarking Oxford Nanopore Read Alignment-Based Structural Variant Detection Tools in Crop Plant Genomes"

### Supplemental Figure 1

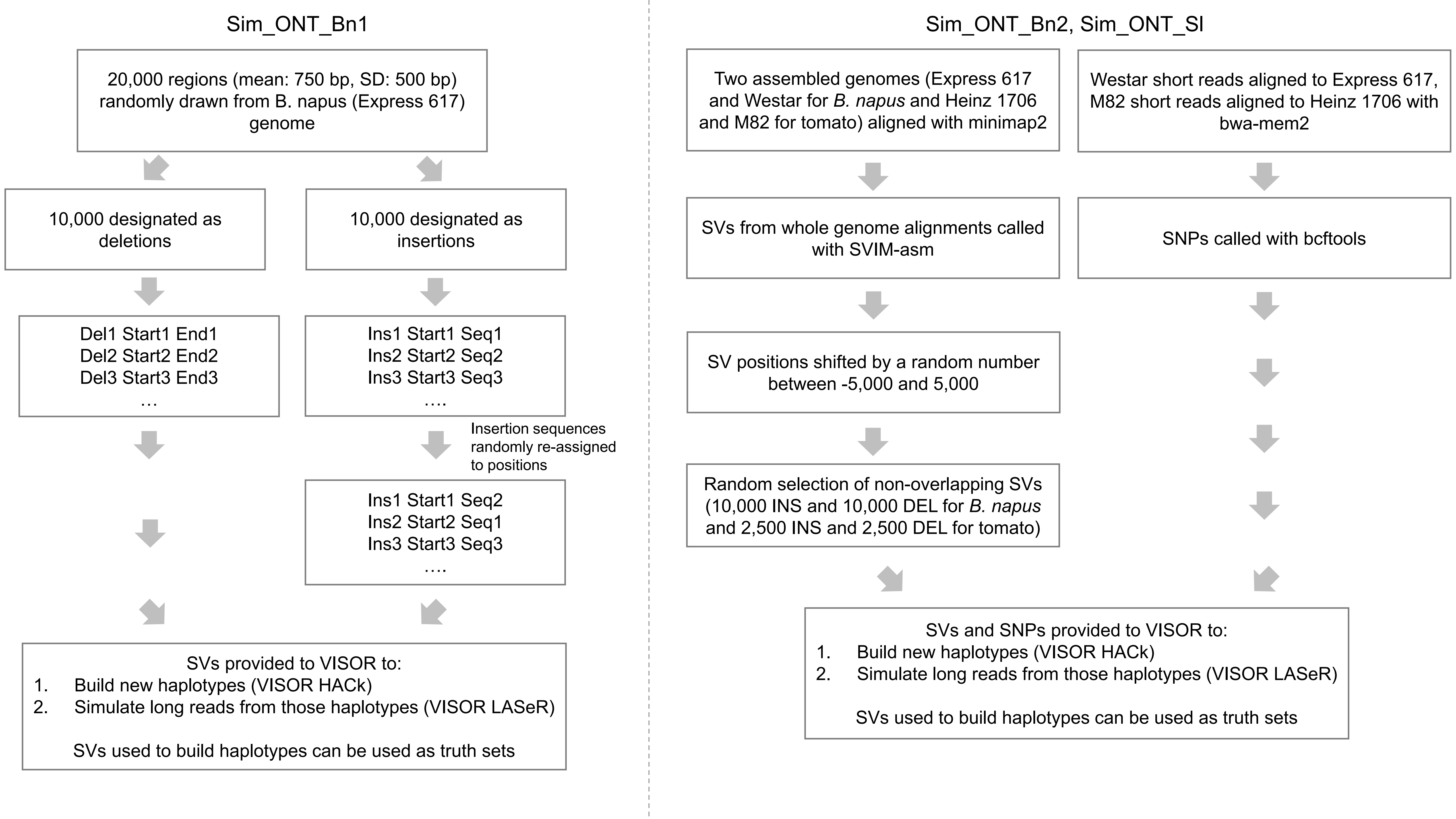
