## Supplemental Tables for "Benchmarking Oxford Nanopore Read Alignment-Based Structural Variant Detection Tools in Crop Plant Genomes"

Table S1: Aligners and SV callers used in the analysis.

| Program Name | Type | Version | Version-Release Year | Citation |
|---|---|---|---|---|
| NGMLR | Aligner | 0.2.7 | 2018 | (Sedlazeck et al., 2018) |
| Vulcan | Aligner | 1.0.3 | 2021 | (Fu et al., 2021) |
| minimap2 | Aligner | 2.24 | 2021 | (Li, 2018, 2021) |
| lra | Aligner | 1.3.2 | 2021 | (Ren and Chaisson, 2021) |
| Sniffles2 | SV caller | 2.0.2 | 2022 | (Sedlazeck et al., 2018) |
| SVIM | SV caller | 2.0.0 | 2021 | (Heller and Vingron, 2019) |
| cuteSV | SV caller | 1.0.13 | 2022 | (Jiang et al., 2020) |
| dysgu | SV caller | 1.3.5 | 2022 | (Cleal and Baird, 2022) |
| NanoVar | SV caller | 1.3.2 | 2020 | (Tham et al., 2019) |

Table S2: Aligner run times and mapping statistics for Real_ONT_Bn, Real_ONT_Sl, Sim_ONT_Bn1, Sim_ONT_Bn2, and Sim_ONT_Sl.

| **Real Datasets** | **Real_ONT_Bn** | | | **Real_ONT_Sl** | | |
|---|---|---|---|---|---|---|
| Aligner | **lra** | | | **lra** | | |
| Coverage | **5x** | **10x** | **20x** | **5x** | **10x** | **20x** |
| Elapsed time (h:mm:ss or m:ss) | 01:13:05 | 2:18:07 | 4:44:12 | 46:34.3 | 01:25:41 | 02:35:05 |
| Mapping statistics | 89.12% | 89.11% | 89.15% | 91.17% | 91.11% | 91.17% |
| Aligner | **minimap2** | | | **minimap2** | | |
| Coverage | **5x** | **10x** | **20x** | **5x** | **10x** | **20x** |
| Elapsed time (h:mm:ss or m:ss) | 59:15.0 | 01:55:55 | 03:52:07 | 21:17.4 | 43:40.8 | 01:19:18 |
| Mapping statistics | 96.32% | 96.32% | 96.33% | 96.69% | 96.66% | 96.68% |
| Aligner | **NGMLR** | | | **NGMLR** | | |
| Coverage | **5x** | **10x** | **20x** | **5x** | **10x** | **20x** |
| Elapsed time (h:mm:ss or m:ss) | 55:06:47 | 103:46:26 | 220:11:56 | 08:52:39 | 19:23:59 | 34:17:16 |
| Mapping statistics | 81.00% | 81.01% | 81.04% | 76.32% | 76.19% | 76.31% |
| Aligner | **Vulcan** | | | **Vulcan** | | |
| Coverage | **5x** | **10x** | **20x** | **5x** | **10x** | **20x** |
| Elapsed time (h:mm:ss or m:ss) | 32:10:56 | 57:15:57 | 119:39:11 | 05:38:56 | 11:33:37 | 21:42:18 |
| Mapping statistics | 97.00% | 96.99% | 96.99% | 98.00% | 98.00% | 97.99% |
| **Simulated Datasets** | **Sim_ONT_Bn1** | | | **Sim_ONT_Bn2** | | |
| Aligner | **lra** | | | **lra** | | |
| Coverage | **5x** | **10x** | **20x** | **5x** | **10x** | **20x** |

| | | | | | | |
|---|---|---|---|---|---|---|
| Elapsed time (h:mm:ss or m:ss) | 36:10.9 | 01:11:49 | 02:23:16 | 36:36.8 | 01:12:28 | 02:24:05 |
| Mapping statistics | 96.67% | 96.66% | 96.67% | 96.69% | 96.71% | 96.69% |
| Aligner | **minimap2** | | | **minimap2** | | |
| Coverage | **5x** | **10x** | **20x** | **5x** | **10x** | **20x** |
| Elapsed time | 19:18.1 | 38:07.1 | 01:16:06 | 19:57.9 | 39:48.7 | 01:18:45 |
| Mapping statistics | 98.65% | 98.63% | 98.65% | 98.83% | 98.83% | 98.81% |
| Aligner | **NGMLR** | | | **NGMLR** | | |
| Coverage | **5x** | **10x** | **20x** | **5x** | **10x** | **20x** |
| Elapsed time (h:mm:ss or m:ss) | 04:18:05 | 09:05:54 | 18:27:09 | 04:34:22 | 08:45:46 | 18:18:12 |
| Mapping statistics | 96.62% | 96.65% | 96.64% | 96.33% | 96.31% | 96.28% |
| Aligner | **Vulcan** | | | **Vulcan** | | |
| Coverage | **5x** | **10x** | **20x** | **5x** | **10x** | **20x** |
| Elapsed time (h:mm:ss or m:ss) | 01:16:23 | 02:32:03 | 5:13:13 | 01:29:29 | 02:56:15 | 06:08:58 |
| Mapping statistics | 99.00% | 99.02% | 99.01% | 99.10% | 99.11% | 99.11% |
| **Simulated Dataset** | **Sim_ONT_Sl** | | | **Sim_ONT_Sl** | | |
| Aligner | **lra** | | | **NGMLR** | | |
| Coverage | **5x** | **10x** | **20x** | **5x** | **10x** | **20x** |
| Elapsed time (h:mm:ss or m:ss) | 42:20.4 | 01:27:02 | 02:48:33 | 02:06:16 | 04:11:44 | 07:43:21 |
| Mapping statistics | 97.58% | 97.57% | 97.60% | 97.10% | 97.13% | 97.10% |
| Aligner | **minimap2** | | | **Vulcan** | | |
| Coverage | **5x** | **10x** | **20x** | **5x** | **10x** | **20x** |
| Elapsed time (h:mm:ss or m:ss) | 14:31.9 | 31:09.0 | 00:54:23 | 00:58:26 | 01:40:02 | 03:47:04 |
| Mapping statistics | 98.56% | 98.57% | 98.57% | 98.96% | 98.99% | 98.97% |

Table S3: Precision, Recall, and F1-score values of SVs for all aligner/SV caller combinations for simulated dataset 1 (Sim_ONT_Bn1).

| | **5X** | | | **10X** | | | **20X** | | |
|---|---|---|---|---|---|---|---|---|---|
| | **minimap2** | | | **minimap2** | | | **minimap2** | | |
| **SVs tools** | **Precision** | **Recall** | **F1** | **Precision** | **Recall** | **F1** | **Precision** | **Recall** | **F1** |
| **cuteSV** | 0.9995 | 0.8191 | 0.9003 | 0.9996 | 0.9377 | 0.9676 | 0.9995 | 0.9914 | 0.9955 |
| **cuteSV-del** | 0.9996 | 0.8307 | 0.9074 | 0.9997 | 0.9482 | 0.9733 | 0.9997 | 0.9925 | 0.9961 |
| **cuteSV-ins** | 0.9994 | 0.8073 | 0.8931 | 0.9995 | 0.9272 | 0.9620 | 0.9994 | 0.9903 | 0.9948 |
| **Sniffles2** | 0.9994 | 0.8068 | 0.8928 | 0.9997 | 0.9298 | 0.9635 | 0.9996 | 0.9901 | 0.9948 |
| **Sniffles-del** | 0.9994 | 0.8247 | 0.9037 | 0.9997 | 0.9465 | 0.9724 | 0.9996 | 0.9931 | 0.9963 |
| **Sniffles-ins** | 0.9995 | 0.7889 | 0.8818 | 0.9997 | 0.9130 | 0.9544 | 0.9997 | 0.9870 | 0.9933 |
| **SVIM.2.0** | 0.9997 | 0.6429 | 0.7825 | 0.9998 | 0.9317 | 0.9645 | 0.9998 | 0.9742 | 0.9869 |
| **SVIM-del** | 0.9997 | 0.6627 | 0.7970 | 0.9998 | 0.9447 | 0.9715 | 0.9998 | 0.9917 | 0.9957 |
| **SVIM-ins** | 0.9997 | 0.6229 | 0.7676 | 0.9998 | 0.9185 | 0.9574 | 0.9999 | 0.9566 | 0.9778 |
| **dysgu** | 0.9984 | 0.7581 | 0.8618 | 0.9982 | 0.8912 | 0.9417 | 0.9987 | 0.9574 | 0.9776 |

| | | | | | | | | | |
|---|---|---|---|---|---|---|---|---|---|
| **dysgu-del** | 0.9990 | 0.8282 | 0.9057 | 0.9985 | 0.9471 | 0.9721 | 0.9985 | 0.9919 | 0.9952 |
| **dysgu-ins** | 0.9975 | 0.6876 | 0.8140 | 0.9978 | 0.8351 | 0.9092 | 0.9989 | 0.9227 | 0.9593 |
| | **lra** | | | **lra** | | | **lra** | | |
| **cuteSV** | 0.9992 | 0.7650 | 0.8665 | 0.9994 | 0.8902 | 0.9417 | 0.9992 | 0.9672 | 0.9829 |
| **cuteSV-del** | 0.9987 | 0.7923 | 0.8836 | 0.9991 | 0.9168 | 0.9562 | 0.9986 | 0.9737 | 0.9860 |
| **cuteSV-ins** | 0.9996 | 0.7375 | 0.8488 | 0.9998 | 0.8635 | 0.9267 | 0.9998 | 0.9606 | 0.9798 |
| **Sniffles2** | 0.9993 | 0.7515 | 0.8578 | 0.9993 | 0.8789 | 0.9352 | 0.9993 | 0.9617 | 0.9801 |
| **Sniffles-del** | 0.9990 | 0.7897 | 0.8821 | 0.9990 | 0.9161 | 0.9557 | 0.9989 | 0.9745 | 0.9865 |
| **Sniffles-ins** | 0.9997 | 0.7130 | 0.8324 | 0.9996 | 0.8415 | 0.9138 | 0.9998 | 0.9488 | 0.9736 |
| **SVIM.2.0** | 0.9991 | 0.5740 | 0.7291 | 0.9994 | 0.8791 | 0.9354 | 0.9994 | 0.9600 | 0.9793 |
| **SVIM-del** | 0.9986 | 0.6260 | 0.7696 | 0.9990 | 0.9171 | 0.9563 | 0.9991 | 0.9728 | 0.9857 |
| **SVIM-ins** | 0.9998 | 0.5217 | 0.6857 | 0.9998 | 0.8409 | 0.9135 | 0.9998 | 0.9472 | 0.9728 |
| **dysgu** | 0.9989 | 0.6124 | 0.7593 | 0.9990 | 0.7734 | 0.8718 | 0.9989 | 0.8439 | 0.9148 |
| **dysgu-del** | 0.9986 | 0.7839 | 0.8783 | 0.9986 | 0.9154 | 0.9552 | 0.9985 | 0.9723 | 0.9852 |
| **dysgu-ins** | 0.9993 | 0.4402 | 0.6112 | 0.9995 | 0.6309 | 0.7735 | 0.9994 | 0.7149 | 0.8336 |
| | **Vulcan** | | | **Vulcan** | | | **Vulcan** | | |
| **cuteSV** | 0.9991 | 0.7390 | 0.8495 | 0.9998 | 0.8618 | 0.9256 | 0.9997 | 0.9517 | 0.9751 |
| **cuteSV-del** | 0.9991 | 0.7716 | 0.8707 | 0.9999 | 0.8942 | 0.9441 | 0.9998 | 0.9654 | 0.9823 |
| **cuteSV-ins** | 0.9990 | 0.7062 | 0.8275 | 0.9996 | 0.8292 | 0.9065 | 0.9997 | 0.9379 | 0.9678 |
| **Sniffles2** | 0.9994 | 0.6670 | 0.8000 | 0.9997 | 0.7838 | 0.8787 | 0.9997 | 0.8984 | 0.9463 |
| **Sniffles-del** | 0.9993 | 0.7461 | 0.8544 | 0.9998 | 0.8734 | 0.9323 | 0.9996 | 0.9574 | 0.9780 |
| **Sniffles-ins** | 0.9995 | 0.5876 | 0.7401 | 0.9997 | 0.6938 | 0.8191 | 0.9998 | 0.8391 | 0.9124 |
| **SVIM.2.0** | 0.9996 | 0.5227 | 0.6864 | 0.9998 | 0.8224 | 0.9024 | 0.9998 | 0.8772 | 0.9345 |
| **SVIM-del** | 0.9995 | 0.5780 | 0.7325 | 0.9998 | 0.8850 | 0.9389 | 0.9998 | 0.9628 | 0.9809 |
| **SVIM-ins** | 0.9998 | 0.4670 | 0.6367 | 0.9999 | 0.7594 | 0.8632 | 0.9997 | 0.7913 | 0.8834 |
| **dysgu** | 0.9984 | 0.5930 | 0.7441 | 0.9985 | 0.7033 | 0.8253 | 0.9984 | 0.7614 | 0.8639 |
| **dysgu-del** | 0.9990 | 0.7698 | 0.8695 | 0.9990 | 0.9026 | 0.9484 | 0.9982 | 0.9652 | 0.9814 |
| **dysgu-ins** | 0.9974 | 0.4155 | 0.5866 | 0.9976 | 0.5031 | 0.6689 | 0.9986 | 0.5568 | 0.7150 |
| | **NGMLR** | | | **NGMLR** | | | **NGMLR** | | |
| **cuteSV** | 0.9978 | 0.6678 | 0.8001 | 0.9987 | 0.7693 | 0.8691 | 0.9987 | 0.8562 | 0.9220 |
| **cuteSV-del** | 0.9965 | 0.7395 | 0.8490 | 0.9981 | 0.8450 | 0.9152 | 0.9977 | 0.9059 | 0.9496 |
| **cuteSV-ins** | 0.9995 | 0.5957 | 0.7465 | 0.9994 | 0.6933 | 0.8187 | 0.9998 | 0.8063 | 0.8927 |
| **Sniffles2** | 0.9982 | 0.4846 | 0.6524 | 0.9983 | 0.5599 | 0.7174 | 0.9986 | 0.6252 | 0.7689 |
| **Sniffles-del** | 0.9976 | 0.6958 | 0.8198 | 0.9978 | 0.8071 | 0.8924 | 0.9982 | 0.8771 | 0.9338 |
| **Sniffles-ins** | 1.0000 | 0.2724 | 0.4282 | 0.9997 | 0.3117 | 0.4753 | 0.9997 | 0.3721 | 0.5424 |
| **SVIM.2.0** | 0.9991 | 0.4596 | 0.6295 | 0.9988 | 0.7392 | 0.8496 | 0.9988 | 0.8157 | 0.8980 |
| **SVIM-del** | 0.9986 | 0.5528 | 0.7116 | 0.9981 | 0.8378 | 0.9110 | 0.9980 | 0.9022 | 0.9477 |
| **SVIM-ins** | 1.0000 | 0.3659 | 0.5358 | 0.9997 | 0.6401 | 0.7805 | 0.9997 | 0.7289 | 0.8431 |
| **dysgu** | 0.9980 | 0.4576 | 0.6275 | 0.9977 | 0.5536 | 0.7120 | 0.9968 | 0.5903 | 0.7415 |
| **dysgu-del** | 0.9978 | 0.7295 | 0.8428 | 0.9976 | 0.8640 | 0.9260 | 0.9964 | 0.9139 | 0.9534 |
| **dysgu-ins** | 0.9989 | 0.1846 | 0.3116 | 0.9979 | 0.2419 | 0.3894 | 0.9981 | 0.2654 | 0.4193 |
| | **5X** | | | **10X** | | | **20X** | | |

| | Precision | Recall | F1 | Precision | Recall | F1 | Precision | Recall | F1 |
|---|---|---|---|---|---|---|---|---|---|
| **NanoVar** | 0.9931 | 0.8145 | 0.8950 | 0.9942 | 0.9268 | 0.9593 | 0.9909 | 0.9789 | 0.9848 |
| **NanoVar-del** | 0.9948 | 0.8238 | 0.9012 | 0.9976 | 0.9394 | 0.9676 | 0.9966 | 0.9860 | 0.9913 |
| **NanoVar-ins** | 0.9914 | 0.8051 | 0.8886 | 0.9907 | 0.9141 | 0.9509 | 0.9852 | 0.9716 | 0.9784 |

Table S4: Total number of the SVs (DEL: deletion; INS: insertion) in Sim_ONT_Bn1.

| | **cuteSV + minimap2** | | | **cuteSV + lra** | | | **cuteSV + Vulcan** | | | **cuteSV + NGMLR** | | |
|---|---|---|---|---|---|---|---|---|---|---|---|---|
| **SV type** | **5x** | **10x** | **20x** | **5x** | **10x** | **20x** | **5x** | **10x** | **20x** | **5x** | **10x** | **20x** |
| **TOTAL** | 16389 | 18762 | 19837 | 15312 | 17814 | 19359 | 14793 | 17239 | 19038 | 13385 | 15406 | 17147 |
| **DEL** | 8327 | 9504 | 9948 | 7949 | 9194 | 9770 | 7738 | 8961 | 9675 | 7437 | 8483 | 9098 |
| **INS** | 8062 | 9258 | 9889 | 7363 | 8620 | 9589 | 7055 | 8278 | 9363 | 5948 | 6923 | 8049 |
| | **Sniffles2+ minimap2** | | | **Sniffles2+ lra** | | | **Sniffles2+ Vulcan** | | | **Sniffles2+ NGMLR** | | |
| **SV type** | **5x** | **10x** | **20x** | **5x** | **10x** | **20x** | **5x** | **10x** | **20x** | **5x** | **10x** | **20x** |
| **TOTAL** | 16146 | 18602 | 19808 | 15044 | 17590 | 19247 | 13370 | 15697 | 17985 | 9784 | 11297 | 12598 |
| **DEL** | 8269 | 9487 | 9955 | 7926 | 9189 | 9776 | 7503 | 8771 | 9609 | 7065 | 8185 | 8883 |
| **INS** | 7877 | 9115 | 9853 | 7118 | 8401 | 9471 | 5867 | 6926 | 8376 | 2719 | 3112 | 3715 |
| | **SVIM+ minimap2** | | | **SVIM + lra** | | | **SVIM + Vulcan** | | | **SVIM + NGMLR** | | |
| **SV type** | **5x** | **10x** | **20x** | **5x** | **10x** | **20x** | **5x** | **10x** | **20x** | **5x** | **10x** | **20x** |
| **TOTAL** | 12861 | 18637 | 19487 | 11490 | 17592 | 19211 | 10457 | 16450 | 17548 | 9199 | 14801 | 16334 |
| **DEL** | 6642 | 9468 | 9939 | 6282 | 9198 | 9756 | 5795 | 8870 | 9649 | 5547 | 8411 | 9058 |
| **INS** | 6219 | 9169 | 9548 | 5208 | 8394 | 9455 | 4662 | 7580 | 7899 | 3652 | 6390 | 7276 |
| | **dysgu + minimap2** | | | **dysgu + lra** | | | **dysgu + Vulcan** | | | **dysgu + NGMLR** | | |
| **SV type** | **5x** | **10x** | **20x** | **5x** | **10x** | **20x** | **5x** | **10x** | **20x** | **5x** | **10x** | **20x** |
| **TOTAL** | 15186 | 17856 | 19173 | 12262 | 15484 | 16896 | 11879 | 14086 | 15253 | 9170 | 11097 | 11844 |
| **DEL** | 8307 | 9504 | 9954 | 7866 | 9185 | 9757 | 7721 | 9053 | 9688 | 7326 | 8678 | 9190 |
| **INS** | 6879 | 8352 | 9219 | 4396 | 6299 | 7139 | 4158 | 5033 | 5565 | 1844 | 2419 | 2654 |

| | **NanoVar** | | |
|---|---|---|---|
| SV type | **5x** | **10x** | **20x** |
| **TOTAL** | 16421 | 18659 | 19780 |
| **DEL** | 8307 | 9441 | 9915 |
| **INS** | 8114 | 9218 | 9865 |

Table S5: Precision, Recall, and F1-score values of SVs for all aligner/SV caller combinations for simulated dataset 2 (Sim_ONT_Bn2).

| | **5X** | | | **10X** | | | **20X** | | |
|---|---|---|---|---|---|---|---|---|---|
| | **minimap2** | | | **minimap2** | | | **minimap2** | | |
| **SVs tools** | **Precision** | **Recall** | **F1** | **Precision** | **Recall** | **F1** | **Precision** | **Recall** | **F1** |
| **cuteSV** | 0.9932 | 0.7754 | 0.8709 | 0.9953 | 0.8729 | 0.9301 | 0.9948 | 0.9328 | 0.9628 |
| **cuteSV-del** | 0.9931 | 0.8329 | 0.9060 | 0.9948 | 0.9294 | 0.9609 | 0.9941 | 0.9713 | 0.9825 |
| **cuteSV-ins** | 0.9934 | 0.7179 | 0.8335 | 0.9959 | 0.8165 | 0.8973 | 0.9955 | 0.8942 | 0.9422 |
| **Sniffles2** | 0.9950 | 0.7555 | 0.8589 | 0.9954 | 0.8521 | 0.9182 | 0.9960 | 0.9164 | 0.9545 |
| **Sniffles-del** | 0.9952 | 0.8233 | 0.9011 | 0.9950 | 0.9236 | 0.9580 | 0.9965 | 0.9693 | 0.9827 |

| | | | | | | | | | |
|---|---|---|---|---|---|---|---|---|---|
| **Sniffles-ins** | 0.9948 | 0.6877 | 0.8132 | 0.9959 | 0.7806 | 0.8752 | 0.9954 | 0.8635 | 0.9248 |
| **SVIM** | 0.9951 | 0.5993 | 0.7481 | 0.9967 | 0.8534 | 0.9195 | 0.9971 | 0.9121 | 0.9527 |
| **SVIM-del** | 0.9956 | 0.6605 | 0.7942 | 0.9962 | 0.9169 | 0.9549 | 0.9967 | 0.9621 | 0.9791 |
| **SVIM-ins** | 0.9945 | 0.5382 | 0.6984 | 0.9972 | 0.7900 | 0.8816 | 0.9976 | 0.8622 | 0.9250 |
| **dysgu** | 0.9902 | 0.7604 | 0.8602 | 0.9890 | 0.8805 | 0.9316 | 0.9880 | 0.9200 | 0.9528 |
| **dysgu-del** | 0.9945 | 0.8166 | 0.8968 | 0.9940 | 0.9239 | 0.9576 | 0.9925 | 0.9592 | 0.9756 |
| **dysgu-ins** | 0.9852 | 0.7043 | 0.8214 | 0.9837 | 0.8372 | 0.9045 | 0.9830 | 0.8809 | 0.9292 |
| | **lra** | | | **lra** | | | **lra** | | |
| **cuteSV** | 0.9935 | 0.6779 | 0.8059 | 0.9960 | 0.7774 | 0.8732 | 0.9959 | 0.8553 | 0.9203 |
| **cuteSV-del** | 0.9930 | 0.7572 | 0.8592 | 0.9949 | 0.8650 | 0.9254 | 0.9949 | 0.9366 | 0.9648 |
| **cuteSV-ins** | 0.9942 | 0.5987 | 0.7474 | 0.9973 | 0.6898 | 0.8155 | 0.9972 | 0.7740 | 0.8715 |
| **Sniffles2** | 0.9941 | 0.6738 | 0.8032 | 0.9959 | 0.7831 | 0.8768 | 0.9954 | 0.8617 | 0.9237 |
| **Sniffles-del** | 0.9947 | 0.7379 | 0.8473 | 0.9963 | 0.8526 | 0.9189 | 0.9968 | 0.9219 | 0.9578 |
| **Sniffles-ins** | 0.9933 | 0.6097 | 0.7556 | 0.9954 | 0.7136 | 0.8313 | 0.9939 | 0.8015 | 0.8874 |
| **SVIM** | 0.9954 | 0.5079 | 0.6726 | 0.9972 | 0.7585 | 0.8616 | 0.9977 | 0.8310 | 0.9068 |
| **SVIM-del** | 0.9945 | 0.5808 | 0.7334 | 0.9965 | 0.8506 | 0.9178 | 0.9970 | 0.9170 | 0.9553 |
| **SVIM-ins** | 0.9966 | 0.4350 | 0.6056 | 0.9981 | 0.6664 | 0.7992 | 0.9987 | 0.7451 | 0.8535 |
| **dysgu** | 0.9915 | 0.5463 | 0.7045 | 0.9919 | 0.7128 | 0.8295 | 0.9922 | 0.7724 | 0.8686 |
| **dysgu-del** | 0.9956 | 0.6863 | 0.8125 | 0.9943 | 0.8364 | 0.9085 | 0.9950 | 0.9026 | 0.9466 |
| **dysgu-ins** | 0.9845 | 0.4063 | 0.5752 | 0.9884 | 0.5891 | 0.7382 | 0.9883 | 0.6420 | 0.7784 |
| | **Vulcan** | | | **Vulcan** | | | **Vulcan** | | |
| **cuteSV** | 0.9932 | 0.6697 | 0.8000 | 0.9936 | 0.7635 | 0.8635 | 0.9948 | 0.8422 | 0.9122 |
| **cuteSV-del** | 0.9927 | 0.7384 | 0.8469 | 0.9923 | 0.8404 | 0.9101 | 0.9938 | 0.9142 | 0.9524 |
| **cuteSV-ins** | 0.9939 | 0.6009 | 0.7490 | 0.9952 | 0.6866 | 0.8126 | 0.9960 | 0.7701 | 0.8686 |
| **Sniffles2** | 0.9944 | 0.6089 | 0.7553 | 0.9947 | 0.7034 | 0.8240 | 0.9955 | 0.7819 | 0.8759 |
| **Sniffles-del** | 0.9949 | 0.6881 | 0.8136 | 0.9945 | 0.7944 | 0.8832 | 0.9957 | 0.8724 | 0.9300 |
| **Sniffles-ins** | 0.9936 | 0.5297 | 0.6910 | 0.9950 | 0.6123 | 0.7581 | 0.9954 | 0.6914 | 0.8160 |
| **SVIM** | 0.9951 | 0.4769 | 0.6448 | 0.9963 | 0.7235 | 0.8382 | 0.9969 | 0.7989 | 0.8870 |
| **SVIM-del** | 0.9952 | 0.5404 | 0.7005 | 0.9964 | 0.8073 | 0.8919 | 0.9973 | 0.8814 | 0.9358 |
| **SVIM-ins** | 0.9949 | 0.4134 | 0.5841 | 0.9961 | 0.6396 | 0.7790 | 0.9965 | 0.7164 | 0.8336 |
| **dysgu** | 0.9929 | 0.5697 | 0.7240 | 0.9920 | 0.7271 | 0.8391 | 0.9923 | 0.7857 | 0.8770 |
| **dysgu-del** | 0.9953 | 0.6397 | 0.7788 | 0.9953 | 0.8072 | 0.8914 | 0.9963 | 0.8737 | 0.9310 |
| **dysgu-ins** | 0.9899 | 0.4997 | 0.6642 | 0.9879 | 0.6470 | 0.7819 | 0.9873 | 0.6976 | 0.8175 |
| | **NGMLR** | | | **NGMLR** | | | **NGMLR** | | |
| **cuteSV** | 0.9885 | 0.6389 | 0.7762 | 0.9906 | 0.7303 | 0.8408 | 0.9909 | 0.8052 | 0.8885 |
| **cuteSV-del** | 0.9853 | 0.7128 | 0.8272 | 0.9874 | 0.8079 | 0.8887 | 0.9873 | 0.8794 | 0.9302 |
| **cuteSV-ins** | 0.9926 | 0.5650 | 0.7201 | 0.9945 | 0.6528 | 0.7882 | 0.9952 | 0.7310 | 0.8429 |
| **Sniffles2** | 0.9896 | 0.5682 | 0.7219 | 0.9916 | 0.6559 | 0.7895 | 0.9919 | 0.7347 | 0.8442 |
| **Sniffles-del** | 0.9886 | 0.6519 | 0.7857 | 0.9905 | 0.7492 | 0.8531 | 0.9904 | 0.8295 | 0.9029 |
| **Sniffles-ins** | 0.9910 | 0.4845 | 0.6509 | 0.9931 | 0.5625 | 0.7182 | 0.9938 | 0.6399 | 0.7785 |
| **SVIM** | 0.9933 | 0.4440 | 0.6137 | 0.9956 | 0.6819 | 0.8095 | 0.9961 | 0.7579 | 0.8608 |
| **SVIM-del** | 0.9918 | 0.5202 | 0.6825 | 0.9958 | 0.7773 | 0.8731 | 0.9960 | 0.8554 | 0.9204 |

| | | | | | | | | | |
|---|---|---|---|---|---|---|---|---|---|
| **SVIM-ins** | 0.9954 | 0.3679 | 0.5372 | 0.9954 | 0.5866 | 0.7382 | 0.9961 | 0.6605 | 0.7943 |
| **dysgu** | 0.9943 | 0.5055 | 0.6703 | 0.9910 | 0.6704 | 0.7998 | 0.9924 | 0.7336 | 0.8436 |
| **dysgu-del** | 0.9952 | 0.6071 | 0.7541 | 0.9948 | 0.7808 | 0.8749 | 0.9965 | 0.8476 | 0.9160 |
| **dysgu-ins** | 0.9929 | 0.4040 | 0.5743 | 0.9857 | 0.5601 | 0.7143 | 0.9868 | 0.6196 | 0.7612 |
| | **5X** | | | **10X** | | | **20X** | | |
| | **Precision** | **Recall** | **F1** | **Precision** | **Recall** | **F1** | **Precision** | **Recall** | **F1** |
| **NanoVar** | 0.9744 | 0.6767 | 0.7987 | 0.9718 | 0.7685 | 0.8583 | 0.9623 | 0.8389 | 0.8964 |
| **NanoVar-del** | 0.9836 | 0.7328 | 0.8399 | 0.9860 | 0.8329 | 0.9030 | 0.9848 | 0.9050 | 0.9432 |
| **NanoVar-ins** | 0.9638 | 0.6205 | 0.7550 | 0.9556 | 0.7041 | 0.8108 | 0.9372 | 0.7727 | 0.8471 |

Table S6: Total number of the SVs (DEL: deletion; INS: insertion) in Sim_ONT_Bn2.

| | **cuteSV + minimap2** | | | **cuteSV + lra** | | | **cuteSV + Vulcan** | | | **cuteSV + NGMLR** | | |
|---|---|---|---|---|---|---|---|---|---|---|---|---|
| **SV type** | **5x** | **10x** | **20x** | **5x** | **10x** | **20x** | **5x** | **10x** | **20x** | **5x** | **10x** | **20x** |
| **TOTAL** | 15605 | 17532 | 18743 | 13638 | 15602 | 17167 | 13476 | 15360 | 16923 | 12918 | 14737 | 16244 |
| **DEL** | 8383 | 9339 | 9767 | 7620 | 8690 | 9410 | 7434 | 8466 | 9196 | 7230 | 8178 | 8904 |
| **INS** | 7222 | 8193 | 8976 | 6018 | 6912 | 7757 | 6042 | 6894 | 7727 | 5688 | 6559 | 7340 |
| | **Sniffles2 + minimap2** | | | **Sniffles2 + lra** | | | **Sniffles2 + Vulcan** | | | **Sniffles2 + NGMLR** | | |
| **SV type** | **5x** | **10x** | **20x** | **5x** | **10x** | **20x** | **5x** | **10x** | **20x** | **5x** | **10x** | **20x** |
| **TOTAL** | 15186 | 17117 | 18396 | 13561 | 15721 | 17310 | 12251 | 14149 | 15713 | 11495 | 13240 | 14827 |
| **DEL** | 8278 | 9283 | 9725 | 7427 | 8557 | 9252 | 6924 | 7998 | 8770 | 6608 | 7580 | 8392 |
| **INS** | 6908 | 7834 | 8671 | 6134 | 7164 | 8058 | 5327 | 6151 | 6943 | 4887 | 5660 | 6435 |
| | **SVIM+ minimap2** | | | **SVIM + lra** | | | **SVIM + Vulcan** | | | **SVIM + NGMLR** | | |
| **SV type** | **5x** | **10x** | **20x** | **5x** | **10x** | **20x** | **5x** | **10x** | **20x** | **5x** | **10x** | **20x** |
| **TOTAL** | 12039 | 17116 | 18286 | 10199 | 15203 | 16648 | 9579 | 14516 | 16019 | 8935 | 13690 | 15209 |
| **DEL** | 6631 | 9200 | 9649 | 5837 | 8531 | 9192 | 5427 | 8099 | 8835 | 5242 | 7801 | 8583 |
| **INS** | 5408 | 7916 | 8637 | 4362 | 6672 | 7456 | 4152 | 6417 | 7184 | 3693 | 5889 | 6626 |
| | **dysgu + minimap2** | | | **dysgu + lra** | | | **dysgu + Vulcan** | | | **dysgu + NGMLR** | | |
| **SV type** | **5x** | **10x** | **20x** | **5x** | **10x** | **20x** | **5x** | **10x** | **20x** | **5x** | **10x** | **20x** |
| **TOTAL** | 15350 | 17795 | 18613 | 11013 | 14363 | 15558 | 11468 | 14650 | 15825 | 10162 | 13522 | 14776 |
| **DEL** | 8206 | 9290 | 9658 | 6889 | 8407 | 9066 | 6423 | 8106 | 8764 | 6096 | 7844 | 8501 |
| **INS** | 7144 | 8505 | 8955 | 4124 | 5956 | 6492 | 5045 | 6544 | 7061 | 4066 | 5678 | 6275 |
| | **NanoVar** | | | | | | | | | | | |
| **SV type** | **5x** | **10x** | **20x** | | | | | | | | | |
| **TOTAL** | 13909 | 15841 | 17484 | | | | | | | | | |
| **DEL** | 7451 | 8446 | 9187 | | | | | | | | | |
| **INS** | 6458 | 7395 | 8297 | | | | | | | | | |

Table S7: Precision, Recall, and F1-score values of SVs for all aligner/SV caller combinations for simulated dataset 3 (Sim_ONT_Sl).

| | **5X** | | | **10X** | | | **20X** | | |
|---|---|---|---|---|---|---|---|---|---|
| | **minimap2** | | | **minimap2** | | | **minimap2** | | |
| | **Precision** | **Recall** | **F1** | **Precision** | **Recall** | **F1** | **Precision** | **Recall** | **F1** |

| | | | | | | | | | |
|---|---|---|---|---|---|---|---|---|---|
| **cuteSV** | 0.9814 | 0.7445 | 0.8467 | 0.9870 | 0.8558 | 0.9167 | 0.9866 | 0.8931 | 0.9375 |
| **cuteSV-del** | 0.9813 | 0.8028 | 0.8831 | 0.9840 | 0.9141 | 0.9477 | 0.9850 | 0.9466 | 0.9654 |
| **cuteSV-ins** | 0.9814 | 0.6856 | 0.8073 | 0.9904 | 0.7970 | 0.8833 | 0.9885 | 0.8391 | 0.9077 |
| **Sniffles2** | 0.9841 | 0.7376 | 0.8432 | 0.9884 | 0.8560 | 0.9174 | 0.9889 | 0.8947 | 0.9394 |
| **Sniffles-del** | 0.9851 | 0.7944 | 0.8795 | 0.9904 | 0.9112 | 0.9492 | 0.9895 | 0.9458 | 0.9671 |
| **Sniffles-ins** | 0.9830 | 0.6803 | 0.8041 | 0.9860 | 0.8002 | 0.8835 | 0.9881 | 0.8432 | 0.9099 |
| **SVIM** | 0.9914 | 0.6057 | 0.7520 | 0.9939 | 0.8536 | 0.9184 | 0.9935 | 0.8879 | 0.9377 |
| **SVIM-del** | 0.9916 | 0.6627 | 0.7944 | 0.9934 | 0.9080 | 0.9488 | 0.9928 | 0.9382 | 0.9647 |
| **SVIM-ins** | 0.9912 | 0.5482 | 0.7060 | 0.9945 | 0.7986 | 0.8858 | 0.9942 | 0.8371 | 0.9089 |
| **dysgu** | 0.9126 | 0.7731 | 0.8371 | 0.9019 | 0.8998 | 0.9008 | 0.8750 | 0.9357 | 0.9043 |
| **dysgu-del** | 0.9117 | 0.8129 | 0.8594 | 0.9028 | 0.9365 | 0.9194 | 0.8798 | 0.9699 | 0.9226 |
| **dysgu-ins** | 0.9136 | 0.7330 | 0.8134 | 0.9010 | 0.8626 | 0.8814 | 0.8698 | 0.9011 | 0.8852 |
| | **lra** | | | **lra** | | | **lra** | | |
| **cuteSV** | 0.9885 | 0.6944 | 0.8158 | 0.9936 | 0.8118 | 0.8936 | 0.9933 | 0.8703 | 0.9278 |
| **cuteSV-del** | 0.9874 | 0.7534 | 0.8547 | 0.9905 | 0.8779 | 0.9308 | 0.9915 | 0.9357 | 0.9628 |
| **cuteSV-ins** | 0.9899 | 0.6349 | 0.7736 | 0.9973 | 0.7451 | 0.8530 | 0.9955 | 0.8043 | 0.8897 |
| **Sniffles2** | 0.9897 | 0.7196 | 0.8334 | 0.9918 | 0.8528 | 0.9170 | 0.9895 | 0.9163 | 0.9515 |
| **Sniffles-del** | 0.9925 | 0.7462 | 0.8519 | 0.9968 | 0.8767 | 0.9329 | 0.9974 | 0.9349 | 0.9652 |
| **Sniffles-ins** | 0.9867 | 0.6929 | 0.8141 | 0.9865 | 0.8286 | 0.9007 | 0.9814 | 0.8975 | 0.9376 |
| **SVIM** | 0.9964 | 0.5585 | 0.7158 | 0.9975 | 0.8124 | 0.8955 | 0.9982 | 0.8731 | 0.9315 |
| **SVIM-del** | 0.9961 | 0.6133 | 0.7591 | 0.9968 | 0.8755 | 0.9322 | 0.9979 | 0.9321 | 0.9639 |
| **SVIM-ins** | 0.9968 | 0.5032 | 0.6688 | 0.9984 | 0.7488 | 0.8558 | 0.9985 | 0.8136 | 0.8966 |
| **dysgu** | 0.9490 | 0.6452 | 0.7682 | 0.9501 | 0.8342 | 0.8884 | 0.9398 | 0.8913 | 0.9149 |
| **dysgu-del** | 0.9472 | 0.6984 | 0.8040 | 0.9463 | 0.8980 | 0.9215 | 0.9356 | 0.9570 | 0.9462 |
| **dysgu-ins** | 0.9511 | 0.5916 | 0.7295 | 0.9548 | 0.7699 | 0.8524 | 0.9448 | 0.8250 | 0.8808 |
| | **Vulcan** | | | **Vulcan** | | | **Vulcan** | | |
| **cuteSV** | 0.9834 | 0.6926 | 0.8128 | 0.9915 | 0.8042 | 0.8881 | 0.9858 | 0.8556 | 0.9161 |
| **cuteSV-del** | 0.9790 | 0.7494 | 0.8490 | 0.9877 | 0.8707 | 0.9255 | 0.9813 | 0.9253 | 0.9525 |
| **cuteSV-ins** | 0.9887 | 0.6353 | 0.7736 | 0.9962 | 0.7370 | 0.8472 | 0.9913 | 0.7853 | 0.8763 |
| **Sniffles2** | 0.9853 | 0.6751 | 0.8012 | 0.9899 | 0.7896 | 0.8785 | 0.9894 | 0.8459 | 0.9120 |
| **Sniffles-del** | 0.9825 | 0.7221 | 0.8324 | 0.9865 | 0.8486 | 0.9123 | 0.9895 | 0.9040 | 0.9448 |
| **Sniffles-ins** | 0.9885 | 0.6276 | 0.7678 | 0.9939 | 0.7301 | 0.8419 | 0.9893 | 0.7873 | 0.8768 |
| **SVIM** | 0.9941 | 0.5395 | 0.6994 | 0.9967 | 0.7884 | 0.8804 | 0.9955 | 0.8449 | 0.9140 |
| **SVIM-del** | 0.9932 | 0.5892 | 0.7396 | 0.9962 | 0.8526 | 0.9188 | 0.9960 | 0.9036 | 0.9476 |
| **SVIM-ins** | 0.9951 | 0.4895 | 0.6562 | 0.9972 | 0.7237 | 0.8387 | 0.9949 | 0.7857 | 0.8780 |
| **dysgu** | 0.9645 | 0.6622 | 0.7852 | 0.9548 | 0.8090 | 0.8759 | 0.9439 | 0.8586 | 0.8992 |
| **dysgu-del** | 0.9649 | 0.6839 | 0.8005 | 0.9576 | 0.8622 | 0.9074 | 0.9469 | 0.9024 | 0.9241 |
| **dysgu-ins** | 0.9640 | 0.6402 | 0.7694 | 0.9515 | 0.7553 | 0.8421 | 0.9406 | 0.8144 | 0.8730 |
| | **NGMLR** | | | **NGMLR** | | | **NGMLR** | | |
| **cuteSV** | 0.9815 | 0.6755 | 0.8002 | 0.9844 | 0.7783 | 0.8693 | 0.9812 | 0.8314 | 0.9001 |
| **cuteSV-del** | 0.9719 | 0.7349 | 0.8370 | 0.9757 | 0.8550 | 0.9114 | 0.9724 | 0.9060 | 0.9380 |
| **cuteSV-ins** | 0.9935 | 0.6155 | 0.7601 | 0.9954 | 0.7010 | 0.8226 | 0.9920 | 0.7561 | 0.8581 |

| | | | | | | | | | |
|---|---|---|---|---|---|---|---|---|---|
| **Sniffles2** | 0.9779 | 0.6612 | 0.7889 | 0.9809 | 0.7681 | 0.8615 | 0.9829 | 0.8326 | 0.9015 |
| **Sniffles-del** | 0.9708 | 0.7064 | 0.8178 | 0.9748 | 0.8229 | 0.8924 | 0.9790 | 0.8779 | 0.9257 |
| **Sniffles-ins** | 0.9864 | 0.6155 | 0.7580 | 0.9882 | 0.7127 | 0.8282 | 0.9873 | 0.7869 | 0.8758 |
| **SVIM** | 0.9916 | 0.5212 | 0.6832 | 0.9968 | 0.7658 | 0.8662 | 0.9954 | 0.8296 | 0.9050 |
| **SVIM-del** | 0.9890 | 0.5759 | 0.7279 | 0.9957 | 0.8390 | 0.9106 | 0.9951 | 0.8928 | 0.9412 |
| **SVIM-ins** | 0.9948 | 0.4660 | 0.6347 | 0.9982 | 0.6921 | 0.8174 | 0.9958 | 0.7658 | 0.8658 |
| **dysgu** | 0.9655 | 0.6315 | 0.7636 | 0.9549 | 0.7866 | 0.8626 | 0.9439 | 0.8384 | 0.8881 |
| **dysgu-del** | 0.9658 | 0.6695 | 0.7908 | 0.9504 | 0.8550 | 0.9002 | 0.9400 | 0.8996 | 0.9194 |
| **dysgu-ins** | 0.9651 | 0.5932 | 0.7348 | 0.9604 | 0.7176 | 0.8214 | 0.9485 | 0.7767 | 0.8541 |
| | **5X** | | | **10X** | | | **20X** | | |
| | **Precision** | **Recall** | **F1** | **Precision** | **Recall** | **F1** | **Precision** | **Recall** | **F1** |
| **NanoVar** | 0.9574 | 0.6170 | 0.7504 | 0.9510 | 0.7051 | 0.8098 | 0.9370 | 0.7138 | 0.8103 |
| **NanoVar-del** | 0.9457 | 0.7699 | 0.8488 | 0.9407 | 0.8799 | 0.9093 | 0.9240 | 0.9225 | 0.9232 |
| **NanoVar-ins** | 0.9777 | 0.4627 | 0.6282 | 0.9688 | 0.5288 | 0.6841 | 0.9620 | 0.5032 | 0.6608 |

Table S8: Total number of the SVs (DEL: deletion; INS: insertion) in Sim_ONT_Sl.

| | **cuteSV + minimap2** | | | **cuteSV + lra** | | | **cuteSV + Vulcan** | | | **cuteSV + NGMLR** | | |
|---|---|---|---|---|---|---|---|---|---|---|---|---|
| SV type | **5x** | **10x** | **20x** | **5x** | **10x** | **20x** | **5x** | **10x** | **20x** | **5x** | **10x** | **20x** |
| **TOTAL** | 3766 | 4305 | 4496 | 3487 | 4054 | 4351 | 3497 | 4028 | 4314 | 3417 | 3928 | 4212 |
| **DEL** | 2042 | 2319 | 2401 | 1904 | 2210 | 2357 | 1911 | 2202 | 2359 | 1888 | 2190 | 2331 |
| **INS** | 1724 | 1986 | 2095 | 1583 | 1844 | 1994 | 1586 | 1826 | 1955 | 1529 | 1738 | 1881 |
| | **Sniffles2.0 + minimap2** | | | **Sniffles2.0 + lra** | | | **Sniffles2.0 + Vulcan** | | | **Sniffles2.0 + NGMLR** | | |
| SV type | **5x** | **10x** | **20x** | **5x** | **10x** | **20x** | **5x** | **10x** | **20x** | **5x** | **10x** | **20x** |
| **TOTAL** | 3723 | 4301 | 4497 | 3612 | 4268 | 4600 | 3402 | 3964 | 4250 | 3361 | 3898 | 4211 |
| **DEL** | 2015 | 2298 | 2390 | 1878 | 2195 | 2343 | 1835 | 2151 | 2286 | 1821 | 2118 | 2243 |
| **INS** | 1708 | 2003 | 2107 | 1734 | 2073 | 2257 | 1567 | 1813 | 1964 | 1540 | 1780 | 1968 |
| | **SVIM+ minimap2** | | | **SVIM + lra** | | | **SVIM + Vulcan** | | | **SVIM + NGMLR** | | |
| SV type | **5x** | **10x** | **20x** | **5x** | **10x** | **20x** | **5x** | **10x** | **20x** | **5x** | **10x** | **20x** |
| **TOTAL** | 3031 | 4261 | 4438 | 2779 | 4039 | 4337 | 2693 | 3926 | 4215 | 2609 | 3812 | 4137 |
| **DEL** | 1666 | 2279 | 2360 | 1533 | 2188 | 2326 | 1479 | 2135 | 2266 | 1453 | 2101 | 2239 |
| **INS** | 1365 | 1982 | 2078 | 1246 | 1851 | 2011 | 1214 | 1791 | 1949 | 1156 | 1711 | 1898 |
| | **dysgu + minimap2** | | | **dysgu + lra** | | | **dysgu + Vulcan** | | | **dysgu + NGMLR** | | |
| SV type | **5x** | **10x** | **20x** | **5x** | **10x** | **20x** | **5x** | **10x** | **20x** | **5x** | **10x** | **20x** |
| **TOTAL** | 4202 | 4948 | 5308 | 3371 | 4353 | 4702 | 3405 | 4206 | 4515 | 3245 | 4087 | 4407 |
| **DEL** | 2222 | 2585 | 2751 | 1836 | 2363 | 2547 | 1766 | 2247 | 2378 | 1728 | 2243 | 2386 |
| **INS** | 1980 | 2363 | 2557 | 1535 | 1990 | 2155 | 1639 | 1959 | 2137 | 1517 | 1844 | 2021 |

| | **NanoVar** | | |
|---|---|---|---|
| **SV type** | **5x** | **10x** | **20x** |
| **TOTAL** | 3212 | 3711 | 3837 |
| **DEL** | 2033 | 2338 | 2499 |
| **INS** | 1179 | 1373 | 1338 |

Table S9: Total number of SVs (DEL: deletion; INS: insertion) in Real_ONT_Bn.

| | **cuteSV + minimap2** | | | **cuteSV + lra** | | | **cuteSV + Vulcan** | | | **cuteSV + NGMLR** | | |
|---|---|---|---|---|---|---|---|---|---|---|---|---|
| **SV type** | **5x** | **10x** | **20x** | **5x** | **10x** | **20x** | **5x** | **10x** | **20x** | **5x** | **10x** | **20x** |
| **TOTAL** | 15962 | 17938 | 20301 | 11367 | 12848 | 14799 | 9214 | 10402 | 12208 | 9334 | 10500 | 12182 |
| **DEL** | 8878 | 9912 | 11016 | 6465 | 7299 | 8316 | 5240 | 5909 | 6872 | 5469 | 6132 | 7034 |
| **INS** | 7084 | 8026 | 9285 | 4902 | 5549 | 6483 | 3974 | 4493 | 5336 | 3865 | 4368 | 5148 |
| | **Sniffles2 + minimap2** | | | **Sniffles2 + lra** | | | **Sniffles2 + Vulcan** | | | **Sniffles2 + NGMLR** | | |
| **SV type** | **5x** | **10x** | **20x** | **5x** | **10x** | **20x** | **5x** | **10x** | **20x** | **5x** | **10x** | **20x** |
| **TOTAL** | 14556 | 16997 | 19070 | 10519 | 12451 | 14401 | 7926 | 9190 | 11006 | 7980 | 9301 | 10895 |
| **DEL** | 8218 | 9501 | 10335 | 5889 | 6943 | 7842 | 4700 | 5430 | 6459 | 4894 | 5636 | 6549 |
| **INS** | 6338 | 7496 | 8735 | 4630 | 5508 | 6559 | 3226 | 3760 | 4547 | 3086 | 3665 | 4346 |
| | **SVIM+ minimap2** | | | **SVIM + lra** | | | **SVIM + Vulcan** | | | **SVIM + NGMLR** | | |
| **SV type** | **5x** | **10x** | **20x** | **5x** | **10x** | **20x** | **5x** | **10x** | **20x** | **5x** | **10x** | **20x** |
| **TOTAL** | 10580 | 15911 | 16906 | 7499 | 11796 | 13203 | 5568 | 9244 | 10596 | 5625 | 9283 | 10596 |
| **DEL** | 6210 | 9163 | 9413 | 4642 | 7229 | 7956 | 3420 | 5659 | 6415 | 3599 | 5842 | 6594 |
| **INS** | 4370 | 6748 | 7493 | 2857 | 4567 | 5247 | 2148 | 3585 | 4181 | 2026 | 3441 | 4002 |
| | **dysgu + minimap2** | | | **dysgu + lra** | | | **dysgu + Vulcan** | | | **dysgu + NGMLR** | | |
| **SV type** | **5x** | **10x** | **20x** | **5x** | **10x** | **20x** | **5x** | **10x** | **20x** | **5x** | **10x** | **20x** |
| **TOTAL** | 13473 | 17890 | 19717 | 7370 | 11640 | 13108 | 5796 | 9701 | 12049 | 4723 | 9024 | 11710 |
| **DEL** | 6989 | 9073 | 9977 | 4250 | 6166 | 6926 | 3099 | 5204 | 6381 | 2909 | 5272 | 6579 |
| **INS** | 6484 | 8817 | 9740 | 3120 | 5474 | 6182 | 2697 | 4497 | 5668 | 1814 | 3752 | 5131 |

| | **NanoVar** | | |
|---|---|---|---|
| **SV type** | **5x** | **10x** | **20x** |
| **TOTAL** | 4690 | 4960 | 5731 |
| **DEL** | 3018 | 3208 | 3793 |
| **INS** | 1672 | 1752 | 1938 |

Table S10: Total number of SVs (DEL: deletion; INS: insertion) in Real_ONT_Sl.

| | **cuteSV + minimap2** | | | **cuteSV + lra** | | | **cuteSV + Vulcan** | | | **cuteSV + NGMLR** | | |
|---|---|---|---|---|---|---|---|---|---|---|---|---|
| **SV type** | **5x** | **10x** | **20x** | **5x** | **10x** | **20x** | **5x** | **10x** | **20x** | **5x** | **10x** | **20x** |
| **TOTAL** | 4176 | 4946 | 5771 | 3625 | 4424 | 5255 | 3565 | 4270 | 5057 | 3606 | 4307 | 5113 |
| **DEL** | 1871 | 2144 | 2483 | 1602 | 1923 | 2268 | 1615 | 1889 | 2214 | 1686 | 1947 | 2290 |
| **INS** | 2305 | 2802 | 3288 | 2023 | 2501 | 2987 | 1950 | 2381 | 2843 | 1920 | 2360 | 2823 |
| | **Sniffles2.0+ minimap2** | | | **Sniffles2.0 + lra** | | | **Sniffles2.0 + Vulcan** | | | **Sniffles2.0 + NGMLR** | | |
| **SV type** | **5x** | **10x** | **20x** | **5x** | **10x** | **20x** | **5x** | **10x** | **20x** | **5x** | **10x** | **20x** |
| **TOTAL** | 4018 | 4937 | 5767 | 3718 | 4641 | 5539 | 3392 | 4223 | 5055 | 3459 | 4278 | 5120 |
| **DEL** | 1764 | 2107 | 2439 | 1548 | 1923 | 2280 | 1524 | 1861 | 2214 | 1595 | 1918 | 2260 |
| **INS** | 2254 | 2830 | 3328 | 2170 | 2718 | 3259 | 1868 | 2362 | 2841 | 1864 | 2360 | 2860 |
| | **SVIM+ minimap2** | | | **SVIM + lra** | | | **SVIM + Vulcan** | | | **SVIM + NGMLR** | | |
| **SV type** | **5x** | **10x** | **20x** | **5x** | **10x** | **20x** | **5x** | **10x** | **20x** | **5x** | **10x** | **20x** |
| **TOTAL** | 2756 | 4701 | 5434 | 2403 | 4267 | 5019 | 2189 | 3955 | 4664 | 2199 | 3952 | 4708 |
| **DEL** | 1250 | 2087 | 2378 | 1115 | 1938 | 2277 | 1015 | 1875 | 2108 | 1063 | 1811 | 2189 |

| INS | 1506 | 2614 | 3056 | 1288 | 2329 | 2742 | 1174 | 2080 | 2556 | 1136 | 2141 | 2519 |
|---|---|---|---|---|---|---|---|---|---|---|---|---|
| | dysgu + minimap2 | | | dysgu + lra | | | dysgu + Vulcan | | | dysgu + NGMLR | | |
| SV type | 5x | 10x | 20x | 5x | 10x | 20x | 5x | 10x | 20x | 5x | 10x | 20x |
| TOTAL | 4643 | 6255 | 7746 | 3278 | 5076 | 6080 | 3129 | 4545 | 5747 | 2999 | 4559 | 5768 |
| DEL | 1992 | 2701 | 3370 | 1387 | 2227 | 2713 | 1302 | 1978 | 2512 | 1269 | 2037 | 2596 |
| INS | 2651 | 3554 | 4376 | 1891 | 2849 | 3367 | 1827 | 2567 | 3235 | 1730 | 2522 | 3172 |

| | NanoVar | | |
|---|---|---|---|
| SV type | 5x | 10x | 20x |
| TOTAL | 1919 | 2209 | 2748 |
| DEL | 1080 | 1219 | 1568 |
| INS | 839 | 990 | 1180 |

Table S11: Proportion of overlapped SVs, deletions and insertions for 5x, 10x and 20x coverages for simulated datasets.

| | Sim_ONT_Bn1 | | | | | Sim_ONT_Bn2 | | | |
|---|---|---|---|---|---|---|---|---|---|
| | All SVs | | | | | All SVs | | | |
| | minimap2 | lra | NGMLR | Vulcan | | minimap2 | lra | NGMLR | Vulcan |
| cuteSV | 76.99% | 69.93% | 62.74% | 66.00% | cuteSV | 73.95% | 65.34% | 63.14% | 64.63% |
| dysgu | 67.57% | 54.81% | 57.16% | 54.95% | dysgu | 73.23% | 56.45% | 55.93% | 60.55% |
| SVIM | 59.38% | 52.43% | 41.89% | 44.37% | SVIM | 57.89% | 51.13% | 47.79% | 49.56% |
| Sniffles2 | 75.35% | 67.97% | 58.11% | 57.09% | Sniffles2 | 72.08% | 63.96% | 59.84% | 61.96% |
| NanoVar | 75.07% | | | | NanoVar | 60.55% | | | |
| | Deletions | | | | | Deletions | | | |
| | minimap2 | lra | NGMLR | Vulcan | | minimap2 | lra | NGMLR | Vulcan |
| cuteSV | 79.19% | 74.45% | 71.30% | 71.28% | cuteSV | 80.05% | 71.07% | 68.41% | 69.40% |
| dysgu | 78.71% | 73.76% | 71.30% | 71.69% | dysgu | 78.95% | 64.70% | 61.66% | 63.55% |
| SVIM | 63.04% | 59.17% | 54.15% | 53.44% | SVIM | 63.90% | 55.46% | 51.98% | 52.84% |
| Sniffles2 | 78.35% | 74.13% | 66.38% | 67.39% | Sniffles2 | 79.14% | 69.71% | 63.86% | 65.60% |
| NanoVar | 77.92% | | | | NanoVar | 66.66% | | | |
| | Insertions | | | | | Insertions | | | |
| | minimap2 | lra | NGMLR | Vulcan | | minimap2 | lra | NGMLR | Vulcan |
| cuteSV | 74.79% | 65.40% | 53.50% | 60.66% | cuteSV | 67.44% | 58.71% | 57.02% | 59.16% |
| dysgu | 56.28% | 33.04% | 22.74% | 32.53% | dysgu | 67.28% | 45.62% | 48.50% | 56.94% |
| SVIM | 55.70% | 45.62% | 28.55% | 34.86% | SVIM | 51.41% | 45.97% | 42.55% | 45.64% |
| Sniffles2 | 72.33% | 61.75% | 40.85% | 46.10% | Sniffles2 | 64.40% | 57.63% | 54.74% | 57.47% |
| NanoVar | 72.26% | | | | NanoVar | 54.20% | | | |

| | Sim_ONT_Sl | | | |
|---|---|---|---|---|
| | All SVs | | | |
| | minimap2 | lra | NGMLR | Vulcan |
| cuteSV | 73.49% | 69.12% | 67.88% | 69.27% |
| dysgu | 68.82% | 60.38% | 63.07% | 65.31% |
| SVIM | 60.62% | 55.74% | 53.59% | 55.44% |

| | | | | |
|---|---|---|---|---|
| **Sniffles2** | 72.73% | 68.89% | 65.18% | 67.46% |
| **NanoVar** | 51.58% | | | |
| | **Deletions** | | | |
| | **minimap2** | **lra** | **NGMLR** | **Vulcan** |
| **cuteSV** | 77.52% | 71.01% | 69.61% | 70.88% |
| **dysgu** | 72.62% | 62.98% | 63.53% | 65.80% |
| **SVIM** | 64.24% | 58.64% | 56.70% | 57.77% |
| **Sniffles2** | 76.32% | 70.85% | 66.40% | 68.69% |
| **NanoVar** | 71.26% | | | |
| | **Insertions** | | | |
| | **minimap2** | **lra** | **NGMLR** | **Vulcan** |
| **cuteSV** | 68.98% | 66.91% | 65.75% | 67.36% |
| **dysgu** | 64.79% | 57.34% | 62.54% | 64.85% |
| **SVIM** | 56.59% | 52.42% | 49.95% | 52.74% |
| **Sniffles2** | 68.72% | 66.84% | 63.77% | 66.01% |
| **NanoVar** | 27.71% | | | |

Table S12: Proportion of overlapped SVs, deletions and insertions for 5x, 10x and 20x coverages for real-world data.

| | **Real_ONT_Sl** | | | | | **Real_ONT_Bn** | | | |
|---|---|---|---|---|---|---|---|---|---|
| | **All SVs** | | | | | **All SVs** | | | |
| | **minimap2** | **lra** | **NGMLR** | **Vulcan** | | **minimap2** | **lra** | **NGMLR** | **Vulcan** |
| **cuteSV** | 49.34% | 47.24% | 47.84% | 47.25% | **cuteSV** | 51.53% | 48.20% | 44.68% | 42.94% |
| **dysgu** | 40.82% | 36.84% | 37.92% | 39.24% | **dysgu** | 46.02% | 35.52% | 26.50% | 30.22% |
| **SVIM** | 34.87% | 33.78% | 31.61% | 32.01% | **SVIM** | 40.01% | 37.71% | 34.79% | 33.44% |
| **Sniffles2** | 47.80% | 46.30% | 45.84% | 45.35% | **Sniffles2** | 50.10% | 45.36% | 42.97% | 40.77% |
| **NanoVar** | 21.57% | | | | **NanoVar** | 19.04% | | | |
| | **Deletions** | | | | | **Deletions** | | | |
| | **minimap2** | **lra** | **NGMLR** | **Vulcan** | | **minimap2** | **lra** | **NGMLR** | **Vulcan** |
| **cuteSV** | 49.63% | 49.16% | 49.72% | 48.86% | **cuteSV** | 54.52% | 51.62% | 47.27% | 44.93% |
| **dysgu** | 40.91% | 36.37% | 37.52% | 38.37% | **dysgu** | 50.71% | 40.53% | 30.48% | 32.21% |
| **SVIM** | 36.04% | 35.89% | 34.77% | 34.19% | **SVIM** | 43.43% | 41.33% | 37.46% | 35.80% |
| **Sniffles2** | 49.41% | 48.52% | 48.00% | 47.36% | **Sniffles2** | 54.56% | 49.88% | 45.49% | 43.00% |
| **NanoVar** | 31.20% | | | | **NanoVar** | 25.07% | | | |
| | **Insertions** | | | | | **Insertions** | | | |
| | **minimap2** | **lra** | **NGMLR** | **Vulcan** | | **minimap2** | **lra** | **NGMLR** | **Vulcan** |
| **cuteSV** | 49.16% | 45.79% | 46.30% | 45.97% | **cuteSV** | 47.91% | 43.98% | 41.21% | 40.43% |
| **dysgu** | 40.73% | 37.19% | 38.24% | 39.87% | **dysgu** | 41.50% | 30.10% | 21.46% | 28.07% |
| **SVIM** | 33.98% | 32.07% | 28.90% | 30.24% | **SVIM** | 35.65% | 32.49% | 30.51% | 29.91% |
| **Sniffles2** | 46.61% | 44.78% | 44.12% | 43.79% | **Sniffles2** | 44.92% | 40.20% | 39.24% | 37.71% |
| **NanoVar** | 12.16% | | | | **NanoVar** | 10.21% | | | |

Table S13: The number of homozygous and heterozygous SVs in simulated datasets (from Sim_ONT_Bn1 to Sim_ONT_Sl).

| | 5x/Sim_ONT_Bn1 | | 10x/Sim_ONT_Bn1 | | 20x/Sim_ONT_Bn1 | |
|---|---|---|---|---|---|---|
| | **Homozygous (1/1)** | **Heterozygous (0/1)** | **Homozygous (1/1)** | **Heterozygous (0/1)** | **Homozygous (1/1)** | **Heterozygous (0/1)** |
| | **minimap2** | | **minimap2** | | **minimap2** | |
| **cuteSV** | 16389 | 37 | 18762 | 48 | 19837 | 51 |
| **Sniffles2** | 16146 | 43 | 18602 | 33 | 19808 | 28 |
| **SVIM** | 12861 | 57 | 18637 | 57 | 19487 | 40 |
| **dysgu** | 15186 | 1276 | 17856 | 1038 | 19173 | 743 |
| | **lra** | | **lra** | | **lra** | |
| **cuteSV** | 15312 | 83 | 17814 | 85 | 19359 | 124 |
| **Sniffles2** | 15044 | 182 | 17590 | 111 | 19247 | 116 |
| **SVIM** | 11490 | 180 | 17592 | 128 | 19211 | 138 |
| **dysgu** | 12262 | 1786 | 15484 | 2235 | 16896 | 2469 |
| | **Vulcan** | | **Vulcan** | | **Vulcan** | |
| **cuteSV** | 14793 | 65 | 17239 | 127 | 19038 | 156 |
| **Sniffles2** | 13370 | 42 | 15697 | 48 | 17985 | 48 |
| **SVIM** | 10457 | 158 | 16450 | 159 | 17548 | 203 |
| **dysgu** | 11879 | 2359 | 14086 | 2822 | 15253 | 3554 |
| | **NGMLR** | | **NGMLR** | | **NGMLR** | |
| **cuteSV** | 13385 | 95 | 15406 | 189 | 17147 | 314 |
| **Sniffles2** | 9784 | 41 | 11297 | 41 | 12598 | 69 |
| **SVIM** | 9199 | 425 | 14801 | 410 | 16334 | 628 |
| **dysgu** | 9170 | 1401 | 11097 | 1965 | 11844 | 2513 |
| | **Homozygous (1/1)** | **Heterozygous (0/1)** | **Homozygous (1/1)** | **Heterozygous (0/1)** | **Homozygous (1/1)** | **Heterozygous (0/1)** |
| **NanoVar** | 16421 | 26 | 18658 | 24 | 19778 | 32 |
| | **5x/Sim_ONT_Bn2** | | **10x/Sim_ONT_Bn2** | | **20x/Sim_ONT_Bn2** | |
| | **Homozygous (1/1)** | **Heterozygous (0/1)** | **Homozygous (1/1)** | **Heterozygous (0/1)** | **Homozygous (1/1)** | **Heterozygous (0/1)** |
| | **minimap2** | | **minimap2** | | **minimap2** | |
| **cuteSV** | 15605 | 507 | 17532 | 607 | 18743 | 812 |
| **Sniffles2** | 15186 | 580 | 17117 | 659 | 18396 | 768 |
| **SVIM** | 12039 | 794 | 17116 | 883 | 18286 | 1047 |
| **dysgu** | 15350 | 922 | 17795 | 953 | 18613 | 1035 |
| | **lra** | | **lra** | | **lra** | |
| **cuteSV** | 13638 | 421 | 15602 | 550 | 17167 | 692 |
| **Sniffles2** | 13561 | 777 | 15721 | 791 | 17310 | 919 |
| **SVIM** | 10199 | 908 | 15203 | 962 | 16648 | 1125 |
| **dysgu** | 11013 | 1118 | 14363 | 1678 | 15558 | 2041 |
| | **Vulcan** | | **Vulcan** | | **Vulcan** | |
| **cuteSV** | 13476 | 324 | 15360 | 444 | 16923 | 559 |
| **Sniffles2** | 12251 | 311 | 14149 | 345 | 15713 | 396 |
| **SVIM** | 9579 | 499 | 14516 | 543 | 16019 | 627 |

| | | | | | | |
|---|---|---|---|---|---|---|
| **dysgu** | 11468 | 609 | 14650 | 652 | 15825 | 696 |
| | **NGMLR** | | **NGMLR** | | **NGMLR** | |
| **cuteSV** | 12918 | 451 | 14737 | 567 | 16244 | 755 |
| **Sniffles2** | 11495 | 302 | 13240 | 348 | 14827 | 413 |
| **SVIM** | 8935 | 540 | 13690 | 584 | 15209 | 737 |
| **dysgu** | 10162 | 541 | 13522 | 665 | 14776 | 672 |
| | **Homozygous (1/1)** | **Heterozygous (0/1)** | **Homozygous (1/1)** | **Heterozygous (0/1)** | **Homozygous (1/1)** | **Heterozygous (0/1)** |
| **NanoVar** | 13908 | 212 | 15841 | 243 | 17483 | 297 |
| | **5x/Sim_ONT_Sl** | | **10x/Sim_ONT_Sl** | | **20x/Sim_ONT_Sl** | |
| | **Homozygous (1/1)** | **Heterozygous (0/1)** | **Homozygous (1/1)** | **Heterozygous (0/1)** | **Homozygous (1/1)** | **Heterozygous (0/1)** |
| | **minimap2** | | **minimap2** | | **minimap2** | |
| **cuteSV** | 3766 | 341 | 4305 | 408 | 4496 | 540 |
| **Sniffles2** | 3723 | 385 | 4301 | 419 | 4497 | 540 |
| **SVIM** | 3031 | 520 | 4261 | 496 | 4438 | 631 |
| **dysgu** | 4202 | 0 | 4948 | 0 | 5308 | 0 |
| | **lra** | | **lra** | | **lra** | |
| **cuteSV** | 3487 | 159 | 4054 | 219 | 4351 | 269 |
| **Sniffles2** | 3612 | 243 | 4268 | 265 | 4600 | 308 |
| **SVIM** | 2779 | 269 | 4039 | 296 | 4337 | 343 |
| **dysgu** | 3371 | 0 | 4353 | 0 | 4702 | 0 |
| | **Vulcan** | | **Vulcan** | | **Vulcan** | |
| **cuteSV** | 3497 | 161 | 4028 | 206 | 4314 | 255 |
| **Sniffles2** | 3402 | 160 | 3964 | 170 | 4250 | 212 |
| **SVIM** | 2693 | 232 | 3926 | 245 | 4215 | 269 |
| **dysgu** | 3405 | 0 | 4206 | 0 | 4515 | 0 |
| | **Ngmlr** | | **Ngmlr** | | **Ngmlr** | |
| **cuteSV** | 3417 | 197 | 3928 | 244 | 4212 | 291 |
| **Sniffles2** | 3361 | 160 | 3898 | 166 | 4211 | 208 |
| **SVIM** | 2609 | 255 | 3812 | 270 | 4137 | 309 |
| **dysgu** | 3245 | 0 | 4087 | 0 | 4407 | 0 |
| | **Homozygous (1/1)** | **Heterozygous (0/1)** | **Homozygous (1/1)** | **Heterozygous (0/1)** | **Homozygous (1/1)** | **Heterozygous (0/1)** |
| **NanoVar** | 3212 | 116 | 3711 | 156 | 3837 | 154 |

Table S14: The number of homozygous and heterozygous SVs in real-world datasets.

| | **5x/*B. napus*** | | **10x/*B. napus*** | | **20x/*B. napus*** | |
|---|---|---|---|---|---|---|
| | **minimap2** | | **minimap2** | | **minimap2** | |
| | **Homozygous (1/1)** | **Heterozygous (0/1)** | **Homozygous (1/1)** | **Heterozygous (0/1)** | **Homozygous (1/1)** | **Heterozygous (0/1)** |
| **cuteSV** | 15962 | 4794 | 17938 | 5746 | 20301 | 9402 |
| **Sniffles2** | 14556 | 5901 | 16997 | 6284 | 19070 | 8148 |
| **SVIM** | 10580 | 7829 | 15911 | 7875 | 16906 | 10140 |

| | | | | | | |
|---|---|---|---|---|---|---|
| **dysgu** | 13473 | 4098 | 17890 | 6134 | 19717 | 8074 |
| | **lra** | | **lra** | | **lra** | |
| **cuteSV** | 11367 | 2671 | 12848 | 3423 | 14799 | 5261 |
| **Sniffles2** | 10519 | 4083 | 12451 | 4527 | 14401 | 5963 |
| **SVIM** | 7499 | 4551 | 11796 | 4904 | 13203 | 6483 |
| **dysgu** | 7370 | 2438 | 11640 | 4655 | 13108 | 6265 |
| | **Vulcan** | | **Vulcan** | | **Vulcan** | |
| **cuteSV** | 9214 | 1807 | 10402 | 2293 | 12208 | 3698 |
| **Sniffles2** | 7926 | 2174 | 9190 | 2397 | 11006 | 3240 |
| **SVIM** | 5568 | 2873 | 9244 | 3094 | 10596 | 4203 |
| **dysgu** | 5796 | 1466 | 9701 | 2513 | 12049 | 3501 |
| | **NGMLR** | | **NGMLR** | | **NGMLR** | |
| **cuteSV** | 9334 | 1597 | 10500 | 2142 | 12182 | 3277 |
| **Sniffles2** | 7980 | 1738 | 9301 | 1918 | 10895 | 2625 |
| **SVIM** | 5625 | 2476 | 9283 | 2702 | 10596 | 3763 |
| **dysgu** | 4723 | 890 | 9024 | 1795 | 11710 | 2562 |
| | **Homozygous (1/1)** | **Heterozygous (0/1)** | **Homozygous (1/1)** | **Heterozygous (0/1)** | **Homozygous (1/1)** | **Heterozygous (0/1)** |
| **NanoVar** | 4690 | 466 | 4960 | 726 | 5731 | 1091 |
| | **5x/*S. lycopersicum*** | | **10x/*S. lycopersicum*** | | **20x/*S. lycopersicum*** | |
| | **minimap2** | | **minimap2** | | **minimap2** | |
| | **Homozygous (1/1)** | **Heterozygous (0/1)** | **Homozygous (1/1)** | **Heterozygous (0/1)** | **Homozygous (1/1)** | **Heterozygous (0/1)** |
| **cuteSV** | 4176 | 910 | 4946 | 1085 | 5771 | 1730 |
| **Sniffles2** | 4018 | 1100 | 4937 | 1163 | 5767 | 1351 |
| **SVIM** | 2756 | 1396 | 4701 | 1304 | 5434 | 1525 |
| **dysgu** | 4643 | 0 | 6255 | 0 | 7746 | 0 |
| | **lra** | | **lra** | | **lra** | |
| **cuteSV** | 3625 | 302 | 4424 | 359 | 5255 | 547 |
| **Sniffles2** | 3718 | 517 | 4641 | 493 | 5539 | 609 |
| **SVIM** | 2403 | 569 | 4267 | 552 | 5019 | 749 |
| **dysgu** | 3278 | 0 | 5076 | 0 | 6080 | 0 |
| | **Vulcan** | | **Vulcan** | | **Vulcan** | |
| **cuteSV** | 3565 | 210 | 4270 | 277 | 5057 | 408 |
| **Sniffles2** | 3392 | 287 | 4223 | 287 | 5055 | 367 |
| **SVIM** | 2189 | 405 | 3952 | 398 | 4664 | 503 |
| **dysgu** | 3129 | 0 | 4545 | 0 | 5747 | 0 |
| | **NGMLR** | | **NGMLR** | | **NGMLR** | |
| **cuteSV** | 3606 | 212 | 4307 | 282 | 5113 | 402 |
| **Sniffles2** | 3459 | 279 | 4278 | 288 | 5120 | 365 |
| **SVIM** | 2199 | 412 | 3955 | 410 | 4708 | 539 |
| **dysgu** | 2999 | 0 | 4559 | 0 | 5768 | 0 |
| | **Homozygous (1/1)** | **Heterozygous (0/1)** | **Homozygous (1/1)** | **Heterozygous (0/1)** | **Homozygous (1/1)** | **Heterozygous (0/1)** |
| **NanoVar** | 1919 | 58 | 2209 | 99 | 2747 | 146 |

Cleal, K. and Baird, D. M. (2022) Dysgu: efficient structural variant calling using short or long reads. *Nucleic acids research* **50**, e53.

Fu, Y., Mahmoud, M., Muraliraman, V. V., Sedlazeck, F. J. and Treangen, T. J. (2021) Vulcan: Improved long-read mapping and structural variant calling via dual-mode alignment. *Gigascience* **10**, giab063.

Heller, D. and Vingron, M. (2019) SVIM: structural variant identification using mapped long reads. *Bioinformatics* **35**, 2907–2915.

Jiang, T., Liu, Y., Jiang, Y., Li, J., Gao, Y., Cui, Z., Liu, Y., Liu, B. and Wang, Y. (2020) Long-read-based human genomic structural variation detection with cuteSV. *Genome Biology* **21**, 189.

Li, H. (2018) Minimap2: pairwise alignment for nucleotide sequences. *Bioinformatics* **34**, 3094–3100.

Li, H. (2021) New strategies to improve minimap2 alignment accuracy. *Bioinformatics* **37**, 4572–4574.

Ren, J. and Chaisson, M. J. P. (2021) lra: A long read aligner for sequences and contigs. *PLOS Computational Biology* **17**, e1009078.

Sedlazeck, F. J., Rescheneder, P., Smolka, M., Fang, H., Nattestad, M., Haeseler, A. von and Schatz, M. C. (2018) Accurate detection of complex structural variations using single-molecule sequencing. *Nat Methods* **15**, 461–468.

Tham, C. Y., Tirado-Magallanes, R., Goh, Y., Fullwood, M. J., Koh, B. T., Wang, W., Ng, C. H., Chng, W. J., Thiery, A., Tenen, D. G. and Benoukraf, T. (2019) NanoVar: Accurate Characterization of Patients' Genomic Structural Variants Using Low-Depth Nanopore Sequencing. *bioRxiv*, 662940.
