## Supplemental Note for "Benchmarking Oxford Nanopore Read Alignment-Based Structural Variant Detection Tools in Crop Plant Genomes"

**Whole genome alignments and SV calling**

minimap2 -ax asm5 --cs -r2k -t 8 target.fa query.fa
svim-asm haploid --min_sv_size 50 --max_sv_size 50000 --types DEL,INS ./wrk reads.bam Ref.fa

**SNP calling**

bwa-mem2 mem -t Ref.fa ${ID}_1.fastq ${ID}_2.fastq
bcftools call --skip-variants indels --threads 1 --ploidy 1 -c -v -Ov -o calls.${ID}.vcf

**Subsampling to 5x, 10x, and 20x coverages**

#for *B. napus*
rasusa –input ${ID}.fq  --coverage 5 --genome-size 1100000000 --output ${ID}.fq
rasusa --input ${ID}.fq  --coverage 10 --genome-size 1100000000 --output ${ID}.fq
rasusa --input ${ID}.fq  --coverage 20 --genome-size 1100000000 --output ${ID}.fq

#for *S. lycopersicum*
rasusa –input ${ID}.fq  --coverage 5 --genome-size 800000000 --output ${ID}.fq
rasusa --input ${ID}.fq  --coverage 10 --genome-size 800000000 --output ${ID}.fq
rasusa --input ${ID}.fq  --coverage 20 --genome-size 800000000 --output ${ID}.fq

**Alignments**

NGMLR
/usr/bin/time –verbose ngmlr -t 8 -r ref.fa -q ${ID}.fq -o ${ID}.ngmlr.sam -x ont --bam-fix
Vulcan
/usr/bin/time --verbose vulcan -i ${ID}.fq -r ref.fa -o ${ID}.vulcan.out -w ${ID}.vulcan.wrk -t 8 -p 80 -ont
Minimap2
minimap2 -d ref.mmi ref.fa # indexing
/usr/bin/time --verbose minimap2 -ax map-ont --MD -t 8 ref.fa ${ID}.fq -o ${ID}.minimap2.sam
Lra
lra index  -ONT ref.fa  # indexing
/usr/bin/time --verbose lra align -ONT ref.fa ${ID}.fq --printMD -t 8 -p s > ${ID}.lra.sam

**SV calling**

Sniffles2
#5x coverage
sniffles --minsupport 3 --threads 8 --input ${ID}.bam --vcf ${ID}.Sniffles2.vcf
#10x coverage
sniffles --minsupport 5 --threads 8 --input ${ID}.bam --vcf ${ID}.Sniffles2.vcf
#20x coverage
sniffles --minsupport 8 --threads 8 --input ${ID}.bam --vcf ${ID}.Sniffles2.vcf

SVIM

```
svim alignment --interspersed_duplications_as_insertions ${ID}.svim ${ID}.bam ref.fa
```

cuteSV

```
#5x coverage
cutesv -t 8 –s 3 –genotype --max_cluster_bias_INS 100 --diff_ratio_merging_INS 0.3 --max_cluster_bias_DEL 100 --diff_ratio_merging_DEL 0.3 ${ID}.bam ref.fa ${ID}.vcf ${ID}.cuteSV.out
#10x coverage
cutesv -t 8 -s 5 –genotype --max_cluster_bias_INS 100 --diff_ratio_merging_INS 0.3 --max_cluster_bias_DEL 100 --diff_ratio_merging_DEL 0.3 ${ID}.bam ref.fa ${ID}.vcf ${ID}.cuteSV.out
#20x coverage
cutesv -t 8 -s 8 –genotype --max_cluster_bias_INS 100 --diff_ratio_merging_INS 0.3 --max_cluster_bias_DEL 100 --diff_ratio_merging_DEL 0.3 ${ID}.bam ref.fa ${ID}.vcf ${ID}.cuteSV.out
```

dysgu

```
#5x coverage
dysgu call --min-support 3 --mode nanopore ref.fa ${ID}.dysgu ${ID}.bam > ${ID}.dysgu.vcf
#10x coverage
dysgu call --min-support 5 --mode nanopore ref.fa ${ID}.dysgu ${ID}.bam > ${ID}.dysgu.vcf
#20x coverage
dysgu call --min-support 8 --mode nanopore ref.fa ${ID}.dysgu ${ID}.bam > ${ID}.dysgu.vcf
```

NanoVar

```
nanovar -x ont -t 8 ${ID}.fq ref.fa ${ID}.NanoVar
```

**Comparisons**

Truvari

```
bcftools sort read.vcf > read.s.vcf
bcftools view read.s.vcf -Oz -o read.s.vcf.gz
bcftools index read.s.vcf.gz
tabix -p vcf read.s.vcf.gz

truvari bench -b truthset.s.vcf.gz -c read.s.vcf.gz -f ref.fa -o read.truvari
```

Surpyvor

```
surpyvor venn --variants read1.vcf read2.vcf read3.vcf  --keepmerged total.vcf --plotout venn.png > total.counts.tsv
surpyvor upset --variants read1.vcf read2.vcf read3.vcf --plotout upset.png
```
